# Optimized design and *in vivo* application of optogenetically functionalized *Drosophila* dopamine receptors

**DOI:** 10.1101/2023.05.20.541404

**Authors:** Fangmin Zhou, Alexandra-Madelaine Tichy, Bibi Nusreen Imambocus, Francisco J. Rodriguez Jimenez, Marco González Martínez, Ishrat Jahan, Margarita Habib, Nina Wilhelmy, Vanessa Bräuler, Tatjana Lömker, Kathrin Sauter, Charlotte Helfrich-Förster, Jan Pielage, Ilona C. Grunwald Kadow, Harald Janovjak, Peter Soba

## Abstract

Neuromodulatory signaling *via* G protein-coupled receptor (GPCRs) plays a pivotal role in regulating neural network function and animal behavior. Recent efforts have led to the development of optogenetic tools to induce G protein-mediated signaling, with the promise of acute and cell type-specific manipulation of neuromodulatory signals. However, designing and deploying optogenetically functionalized GPCRs (optoXRs) with accurate specificity and activity to mimic endogenous signaling *in vivo* remains challenging. Here we optimized the design of optoXRs by considering evolutionary conserved GPCR-G protein interactions and demonstrate the feasibility of this approach using two *Drosophila* Dopamine receptors (optoDopRs). We validated these optoDopRs showing that they exhibit high signaling specificity and light sensitivity *in vitro*. *In vivo* we detected receptor and cell type-specific effects of dopaminergic signaling in various behaviors including the ability of optoDopRs to rescue loss of the endogenous receptors. This work demonstrates that OptoXRs can enable optical control of neuromodulatory receptor specific signaling in functional and behavioral studies.

## Introduction

Behavioral flexibility, learning as well as goal-directed and state-dependent behavior in animals depend to a large degree on neuromodulatory signaling *via* G protein coupled receptors (GPCRs), which tune neuronal network function to the current external sensory environment and the internal state of the animal ^1^. Dopamine (DA) is one of the most conserved neurotransmitters and modulators, which can activate different G protein-dependent and -independent signaling events ^2, 3^. DA signaling can increase or decrease excitability of the affected neuronal substrates as well as induce synaptic plasticity and long-term transcriptional changes *via* distinct cognate receptors. Through its receptors, DA regulates numerous functional processes including motivation, locomotion, learning and memory ^2–6^. While it is desirable to get more precise insight into the action of DA signaling and other neuromodulators, pharmacological approaches lack the precision and specificity to target defined circuits and their regulated behaviors. At the same time, most current genetic tools lack the temporal control and sensitivity required to manipulate the corresponding receptors directly and acutely with high efficiency *in vivo*.

Optogenetics has revolutionized our understanding of the function of specific neural circuits, allowing for investigation of their role in behavior and physiology through genetic targeting and high spatiotemporal precision ^7–9^. While cell type-specific manipulation of neurons *in vivo* using light-controlled ion channels has evolved rapidly, and numerous powerful tools are available, optical control of modulatory GPCR mediated-signaling in general, and in circuits endogenous to the modulatory neurotransmitter, has been more limited so far ^10–12^. This is in part due to the difficulty of designing functional light-activatable GPCRs showing endogenous-like localization and activity of the target receptor. Previous studies established chimeric receptor designs in which the intracellular domains of a receptor of interest were swapped into a prototypical light sensitive GPCR, typically bovine rhodopsin (Rho). In one example, this strategy has been successfully applied to the β2-adrenergic receptor (β2AR) and has yielded a functional optoXR displaying signaling comparable to its native counterpart^13–17^. A systematic approach for class A GPCRs has produced a library of human optoXRs displaying *in vitro* signaling capacity corresponding to orphan receptors ^18^. Similarly, functional class A/F chimera (Rho:Frizzled7) and class A/C chimera (mOpn4:mGluR6) were designed and applied in optogenetic cellular migration and vision restoration studies, respectively^19, 20^. Additional approaches have used structure-guided design, primary sequence-based empirical methods or native light-sensitive GPCRs with similar signaling properties as the receptor of interest ^10, 11, 17^. While it is appealing to utilize optoXRs to mimic GPCR function, design and functionality remain challenging. Importantly the signaling properties of many GPCRs depend on the cell type, receptor localization and activation kinetics as well as the functional context ^11, 21–24^. Only in few cases have optoXRs been deployed *in vivo* and they have so far mostly been used to acutely manipulate specific G protein signaling pathways (see Table S1). Thus, there is very limited evidence that optoXRs can functionally replace endogenous GPCR function in target tissues.

*In vivo* models including *Drosophila melanogaster* have contributed extensively to our understanding of neuromodulatory GPCR signaling in neural circuit function and behavior ^1, 25–29^. In particular, DA and its receptors have been long studied in *Drosophila* regarding their role in learning, memory and goal directed behaviors ^3, 5, 6, 30–33^. *Drosophila* encodes 4 Dopamine receptors: two D1-like receptors (Dop1R1 and Dop1R2), a D2-like receptor (Dop2R) and Dopamine-Ecdysteroid receptor (DopEcR). Dop1R1 and Dop1R2 display conserved functions in learning and memory by inducing cAMP signaling and intracellular calcium store release, respectively ^31, 34–40^. Yet so far, most acute (i.e. dynamic and short term) cell type specific functions of these receptors, such as the timing and duration of their signaling, could not be manipulated due to the lack of suitable tools. OptoXRs that can be readily studied *in vivo* and allow precise spatiotemporal dissection of endogenous-like dopaminergic signaling and function would solve these issues but are currently not available.

Here, we generated and optimized chimeric optoXRs of *Drosophila melanogaster* Dop1R1 and Dop1R2 by taking advantage of evolutionary constraints of G protein coupling specificity.

We characterized optoDopR signaling *in vitro* and found that our optimized design resulted in improved signaling specificity and light-dependent G protein activation. *In vivo*, expression and subcellular localization were strongly improved, more closely resembling the endogenous receptor distribution. We then demonstrated that optoDopRs *in vivo* can replace or mimic dopamine receptor functionality in various DA-dependent behaviors including locomotion, arousal, learning and operant feeding behavior. Intriguingly, we found cell type and receptor-specific functions using our optoDopRs in innate and adaptive behaviors showing their utility to study DA-dependent function and behavior with high spatiotemporal precision and specificity.

## Results

### Optimization of sequence-based design for optoDopRs

Previous studies have developed sequence- ^14, 20, 41^ or structure-based ^17^ rules for exchanging regions of GPCRs to generate various chimera that display functional signaling of the target receptor yet altered ligand/sensor specificity. Most optoXRs developed so far built on bovine Rhodopsin (Rho) as a light sensitive backbone, mainly due to its well-described structure and function, together with sequence-based rules developed by Kim *et al*. ^14, 16, 18, 42^; these receptors have then been applied *in vitro* and *in vivo* (summarized in Table S1). In the original design rules, transmembrane (TM) helices and intracellular loop (ICL) regions were exchanged. This resulted in chimeric receptors in which all three ICLs with proximal TM residues and the C-terminus of Rho were substituted by the corresponding regions of the target receptor. When we applied this methodology (here termed ‘V1’, Fig. 1A) to *Drosophila* DopRs and six other GPCRs, we failed to produce functional chimera with the exception of Dop1R1 (see Fig. 1C, and data not shown). Therefore, we revised the receptor design based on recently computed evolutionary constraints of G-protein binding to receptors^43^. It became evident that ICL1 was generally not contributing major G-protein binding contacts so we reasoned that retaining Rho ICL1 should not limit signaling but may increase the structural integrity of a chimeric optoXR. In addition, we readjusted the TM7/C-terminus exchange site to accommodate additional G-protein contact sites. These sites have been defined in the evolutionary analysis of GPCR-G protein interactions through inspection of multiple GPCR-G-protein complex structures of class A receptors. Using this approach (termed ‘V2’), we redesigned the optoDop1R1 chimera and studied the effects of these changes (Fig. 1A).

**Figure 1:**
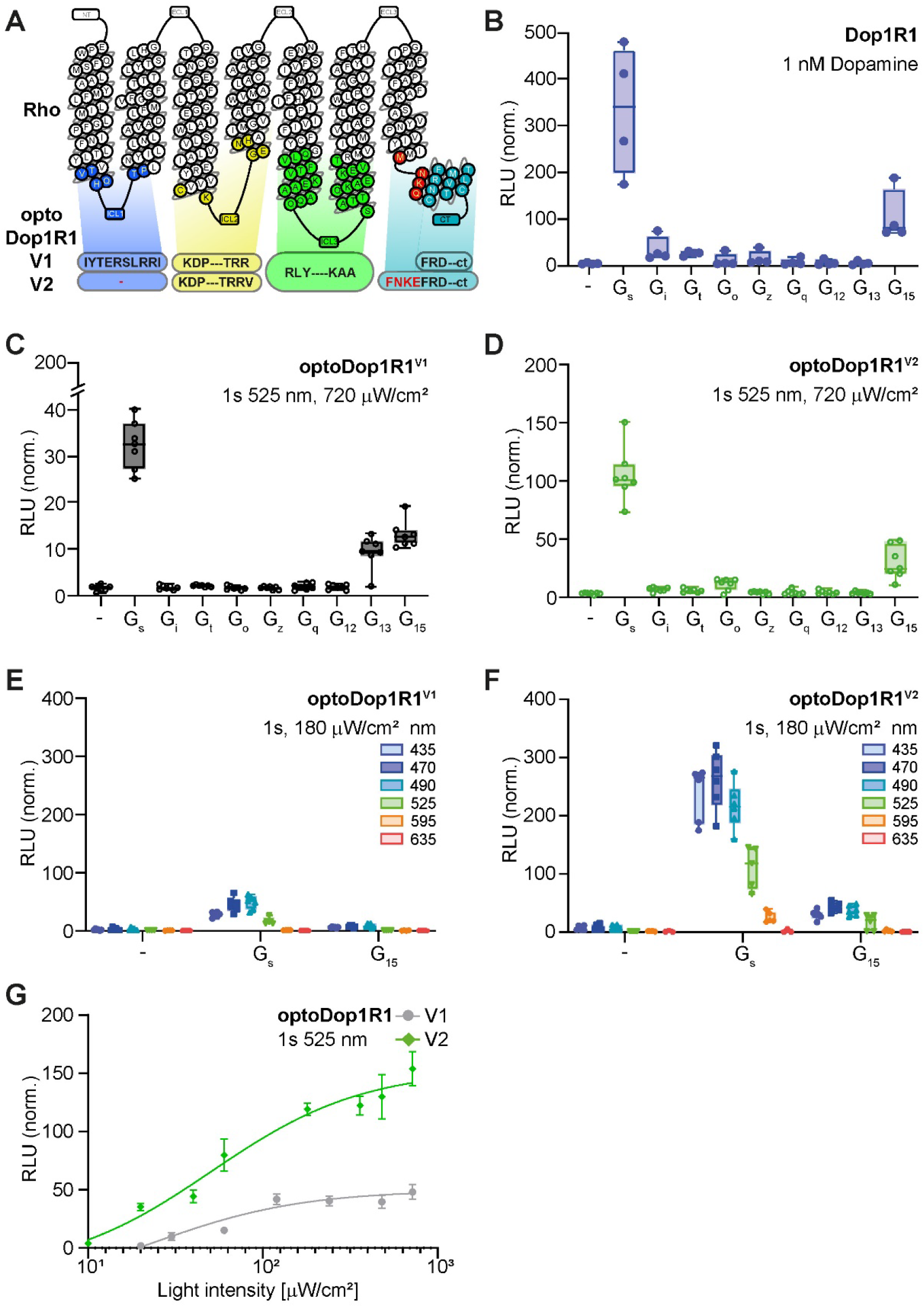
Design and characterization of optoDop1R1^V2^. (A) Schematic overview of optoDop1R1 variants based on the original approach ^14^ (V1) and the optimized design (V2). (B) G protein coupling properties of Drosophila Dop1R1 with 1nM dopamine. Maximum normalized responses are shown (n=4). (C) G protein coupling properties of optoDop1R1^V1^ after activation with light (1s 525 nm, 720 μW/cm^2^). Maximum normalized responses are shown (n=7). (D) G protein coupling properties of improved optoDop1R1^V2^ after activation with light (1s 525 nm, 720 μW/cm^2^). Maximum normalized responses are shown (n=7). (E) Wavelength-dependent maximum G protein activation of optoDop1R1^V1^ after activation with light (1s 180 μW/cm^2^, n=7). (F) Wavelength- dependent maximum G protein coupling of optoDop1R1^V2^ after activation with light (1s 180 μW/cm^2^, n=6). (G) Light intensity-dependent maximum of cAMP induction (G_s_ coupling) of optoDop1R1^V^^1^ and optoDop1R1^V2^ after activation with light (1s 525 nm, n=3-8).

### Characterization of Dop1R1 and optoDop1R1 activation profiles

We compared the activity of the *Drosophila* Dop1R1 receptor with its opto-variants designed with the previous (V1) or optimized (V2) approach. To this end, we utilized a chimeric G_αs_ protein (‘G_sx_‘) assay allowing direct comparison of the coupling and kinetics of GPCRs with the major G-proteins upstream of the cAMP reporter GloSensor ^44^. Upon addition of dopamine, Dop1R1 showed strong coupling to G_s_ as previously described ^39^, as well as G_15_, and moderate coupling to inhibitory G proteins (Fig. 1B, S1A,B). G_s_ and G_15_ coupling showed dose-dependent responses in the pico- to nanomolar range, respectively (Fig. S1B). In comparison, optoDop1R1^V1^ activation using a 1 s light pulse (525 nm) resulted in G_s_, G_13_ and G_15_ coupling with moderate efficiency (Fig. 1C, S1C). While significant activation of G_s_ signaling was observed, the coupling profile did not match the Dop1R1 receptor profile entirely due to aberrant G_13_ signaling and limited coupling to inhibitory G-proteins. In contrast, optoDop1R1^V2^ activation more closely resembled the wildtype receptor displaying strong coupling to G_s_ and G_15_ as well as weak coupling to inhibitory G proteins (Fig. 1D, S1D). We then compared the wavelength dependent G_s_ and G_15_ activation profiles of the two optoDopR variants. While maximum activation was observed with 470-490 nm light, optoDop1R1^V2^ induced 5-10-fold higher responses than the corresponding V1 receptor (Fig. 1E,F, S1E). For optoDop1R1^V2^, significant G_s_ activation was also observed in the green to orange wavelength range up to 595 nm, while optoDop1R1^V1^ activation was virtually undetectable at 595 nm. Direct comparison of light intensity dependent G_s_ signaling induced by V1 vs. V2 showed half-maximal activation at around 50 μW/cm^2^ (at 525 nm) for both optoXRs (Fig. 1G).

However, the V2 design excelled in light sensitivity displaying 3- to 20-fold higher G_s_ responses, particularly at low light intensities below 40 μW/cm^2^. Overall, unlike the classic chimeric sequence-based approach, our optimized optoXR^V2^ design yielded an optoDop1R1 variant exhibiting superior light sensitivity and high signaling specificity comparable to the Dop1R1 wildtype receptor.

### Generation and characterization of functional optoDop1R2^V^^2^

While for Dop1R1 both designs yielded functional optoXRs albeit with different quality, the original approach did not produce a functional optoDop1R2 as no light-dependent responses could be detected in the G_sx_ assay (data not shown). We thus again turned to our optimized design and generated optoDop1R2^V2^, which concordantly contained the Rho ICL1 and the extended C-terminus (Fig. 2A).

**Figure 2:**
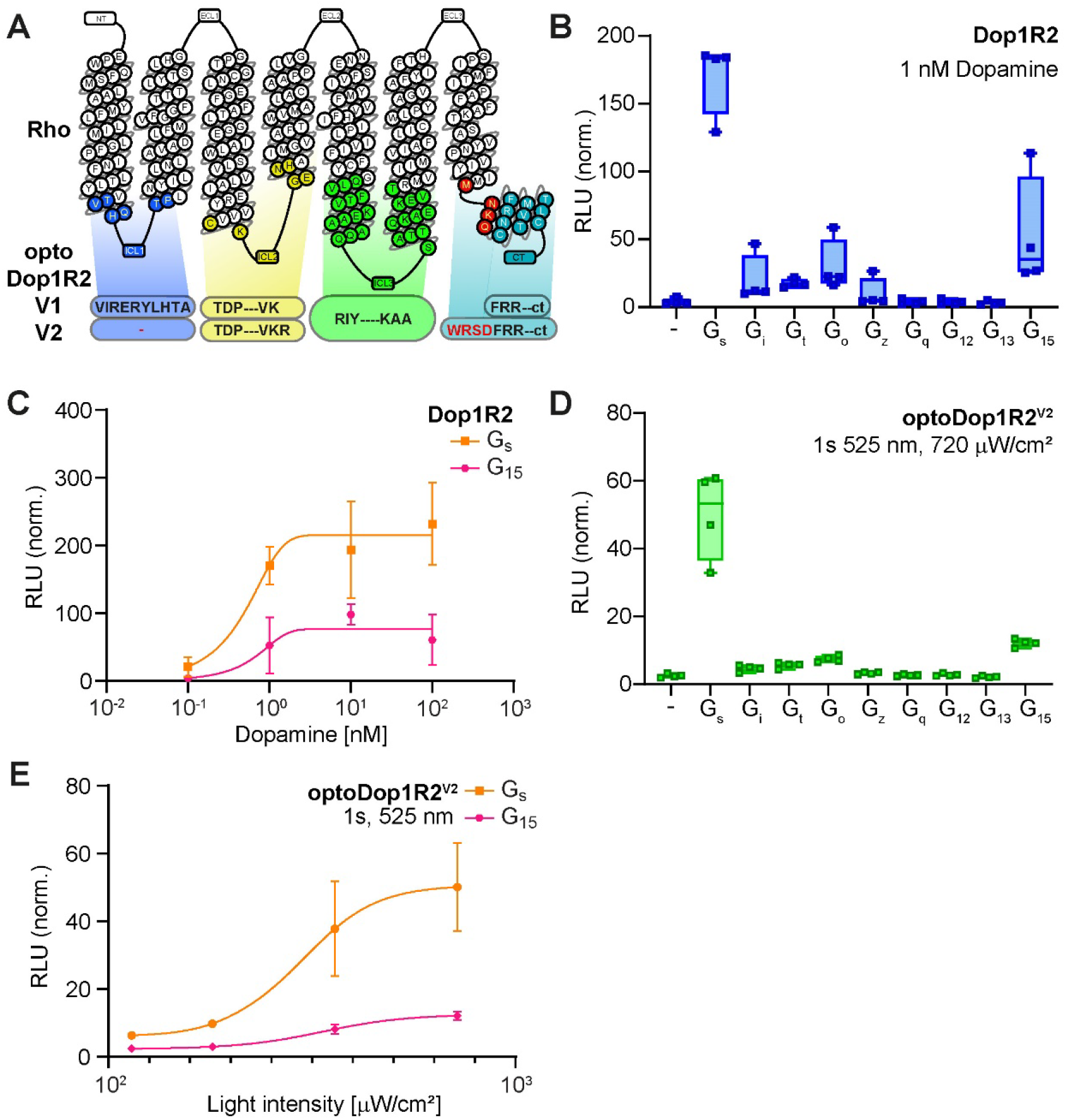
Design and characterization of optoDop1R2^V2^. (A) Schematic overview of optoDop1R2^V2^ design compared to V1. (B) G protein coupling properties of Drosophila Dop1R2 with 1nM dopamine. Maximum normalized responses are shown (n=3-4). (C) DA concentration dependent maximum activation of G_s_ and G_15_ signaling of Dop1R2 (n=3-4). (D) G protein coupling properties of optoDop1R2^V2^ after activation with light (1s 525 nm, 720 μW/cm^2^). Maximum normalized responses are shown (n=4). (E) Light intensity-dependent maximum of G_s_ and G_15_ signaling induced by optoDop1R2^V^^2^ (1s 525 nm, n=4).

We first characterized *Drosophila* Dop1R2 using the G_sx_ assay. Dop1R2 showed dose-dependent coupling to G_s_, G_15_ and inhibitory G proteins upon addition of dopamine in the range of 0.1-100 nM (Fig. 2B,C, S2A,B). For our optimized optoDop1R2^V2^ the implemented changes were indeed sufficient to produce a functional optoXR (Fig. 2D,E, S2C). Similarly to the wildtype receptor, optoDop1R2^V2^ coupled to the same G proteins, prominently with G_s_ and G_15_ showing light dose-dependent responses in the range of 114-720 μW/cm^2^ (Fig. 2D,E). A similar light-dependent profile was obtained for G_i_ and G_o_ responses (Fig. S2D).

The G protein coupling profile and dose-dependent activity of optoDop1R2^V2^ closely resembled the wildtype receptor in this assay, yet the maximum activation levels remained consistently lower under these conditions. As for optoDop1R1^V2^, the rhodopsin-based optoDop1R2^V2^ showed maximum responses to 470-490 nm light (Fig. S2E and data not shown). Overall, these results show that the optoXR^V2^ design approach allowed the generation of functional and specific optoDopRs not obtainable with the previous strategy.

### Characterization of optoDopR localization and functionality in vivo

Based on the promising activity of optoDopRs^V2^ in cell culture assays we generated transgenes to investigate their functionality *in vivo.* We first tested the expression and localization of optoDopRs in the *Drosophila* mushroom body (MB), the central learning and memory center in insects ^45–48^. The principal MB neurons, Kenyon cells (KCs), receive olfactory and other sensory input as well as compartmentalized dopaminergic innervation along their axonal arbors ^48–51^. DopRs are central to MB function and both Dop1R1 and Dop1R2 are involved in learning and memory ^34, 37, 39^. We expressed our optoDopRs in KCs and investigated localization in larval and adult fly MBs. In larval KCs, optoDop1R1^V1^ was detectable in the soma and only weakly in axons (Fig. 3A). Similarly, expression in adult MBs was weak and displayed an irregular pattern (Fig. 3B). In comparison, optoDop1R1^V2^ showed more prominent expression and axonal localization was clearly visible in larval and adult KC axons (Fig. 3A,B). Similarly, optoDop1R2^V2^ showed prominent axonal localization in larval and adult KCs (Fig. 3A,B). We further labeled single Kenyon cells expressing optoDopRs using an activity-induced expression system ^52^. We detected optoDopR expression in the KC soma as well as in punctuated structures along the axon (Fig. S3A,B). Unlike optoDop1R1^V1^, the expression and localization pattern of optoDopRs^V2^ was comparable to recent data of endogenously tagged DopRs ^53^. Overall, these data suggest that the V2 design yielded optoDopRs that more closely resemble endogenous receptor localization with prominent localization along KC axons.

**Figure 3:**
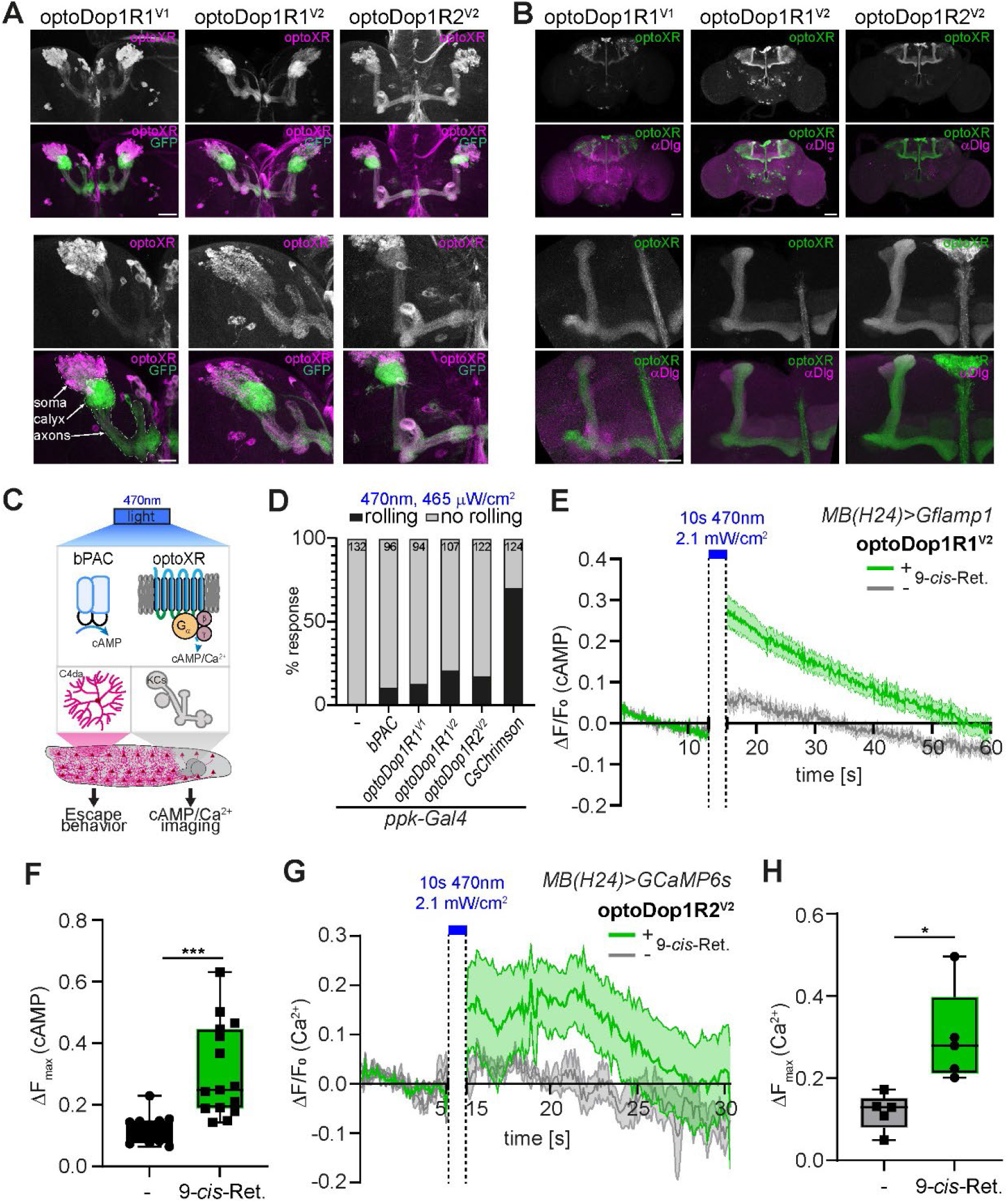
*In vivo* localization and characterization of optoDopR activity. (A) Expression of optoXRs in the larval mushroom body (scale bars: top panels 50 μm, bottom panels 25 μm). (B) Expression of optoXRs in the adult mushroom body (all KCs, *OK107-Gal4*) (scale bars: top panels 50 μm, bottom panels 25 μm). (C) Schematic of blue light induced activation of bPAC and optoXRs expressed in larval nociceptive neurons (C4da) or Kenyon cells (KCs). cAMP increase in C4da neurons elicits spontaneous larval escape responses. KC expression was used to image cAMP and Ca^2+^ responses after bPAC or optoDopR activation. (D) Spontaneous escape responses (rolling) in larvae expressing bPAC, optoXRs or CsChrimson during blue light illumination (470 nm, 465 μW/cm^2^, n as indicated). (E) cAMP imaging in the larval mushroom body using G-Flamp1 and optoDop1R1^V2^ expression (*H24-Gal4>G-Flamp1, optoDop1R1^V^*^2^, 10s 470 nm, n=11, 15). Responses in the medial lobe after 10s blue light illumination are shown over time. (F) Maximum cAMP responses in the MB medial lobe after light-induced activation of optoDop1R1^V2^ (10s 470 nm, n=11,15). (G) *In vivo* calcium imaging of optoDop1R2^V2^ expressed in the larval mushroom body using GCaMP6s before and after activation with light. Neuronal calcium responses in the medial lobe of animals reared with or without 9-*cis*-retinal are shown over time (*H24-Gal4>GCaMP6s, optoDop1R2^V^*^2^, 10s 470 nm, n=5,5). (H) Maximum responses in the MB medial lobe after light-induced activation of optoDop1R2^V2^ (10s 470 nm, n=5,5).

We next wanted to assay if 2^nd^ messenger responses can be elicited by our optoDopRs *in vivo*. Dop1R1 has been reported to be primarily linked to G_s_-dependent cAMP production, while Dop1R2 can induce intracellular calcium release *via* activation of G_q_-family signaling that includes G_15_ ^37, 39^. Elevated cAMP and calcium levels in *Drosophila* larval nociceptors can elicit a stereotyped escape response ^54^, which we chose as a first proxy for functional activation of our optoXRs. We expressed optoDopRs in larval nociceptors and illuminated freely crawling larvae with blue light for 3 min. We expressed the blue light activated adenylate cyclase bPAC ^55^ and the cation channelrhodopsin CsChrimson ^56^ as positive controls for cAMP and calcium-induced escape responses, respectively. bPAC and our optoXRs induced spontaneous rolling during light illumination, which generally occurred sporadically and with some delay (Fig. 3D, movie S1-S4). In contrast, activation of CsChrimson resulted in a high percentage of animals rolling immediately after light onset (movie S5). Consistent with the predicted coupling to intracellular calcium stores by optoDop1R2, we also observed fast rolling responses in some cases. Overall, these data indicate that all optoXRs are capable of inducing 2^nd^ messenger signaling *in vivo* with similarity to cAMP and calcium-induced escape responses.

To measure specific 2^nd^ messenger responses induced by optoDopRs *in vivo*, we used fluorescent reporters for cAMP and calcium levels. Dop1R1 and Dop1R2 were previously shown to primarily regulate cAMP or store-released calcium levels in KC neurons, respectively ^37^. We first expressed the cAMP reporter Gflamp1 ^57^ together with optoDop1R1^V2^ or bPAC in the larval MB and imaged light-induced cAMP changes in the soma and medial lobe regions in dissected live larval brains. bPAC activation with blue light was able to elicit strong cAMP increase particularly in the MB soma region due to its cytosolic localization, and to a lesser extent also in the medial lobe region (Fig. S3C,D, movie S6). In comparison, activation of optoDop1R1^V2^ resulted in cAMP increase preferentially in the medial lobe and to a lower degree in the soma region and was largely dependent on the presence of 9-*cis*- retinal during rearing of the animals (Fig. 3E,F, S3E,F, movie S7). Axonal cAMP levels in the medial lobe decayed to background levels within approx. 60s after a 10s blue light stimulus and could be elicited repeatedly (data not shown). Of note, bPAC has been described to exhibit dark activity ^58^ and baseline fluorescence levels of Gflamp1 were significantly higher than for optoDop1R1^V2^, suggesting optoDop1R1^V2^ exhibits no or low dark activity compared to bPAC.

To test for calcium store release by optoDop1R2^V2^ activation, we expressed it together with the fluorescent calcium reporter GCaMP6s ^59^ in the larval MB and imaged light-induced changes in calcium levels in live intact larvae. Upon blue light illumination, we could detect calcium responses in the medial lobe as well as in KC somata (Fig. 3G,H, S3G,H).

Interestingly, calcium levels remained elevated for up to 10s after light stimulation, similar to store-release of calcium linked to dopaminergic activation in mammalian neurons ^60^. Axonal responses in the medial lobe were overall stronger and more sustained than in the KC somata (Fig. S3G,H) suggesting the local environment of receptor localization affects signaling efficiency.

Taken together, these data show that optoDopRs^V^^2^ display endogenous-like localization and signaling *in vivo*.

### Functional analysis of dopaminergic signaling in fly larvae

We next wanted to test functionality of the optoDopRs in relevant behaviors. Dopamine signaling plays a pivotal and conserved role in locomotion, reward, and innate preference behavior ^2, 3, 5, 31, 61, 62^. Disruption of dopaminergic neuron function in flies and mammals results in locomotion defects and is a key feature of Parkinson’s disease ^63–66^. We used Rotenone- induced impairment of dopaminergic neurons in larvae ^67^, which resulted in reduced locomotion velocity and increased turning behavior as previously described (Fig. 4A). We reasoned that locomotion deficits might be rescued by triggering dopaminergic signaling in the receiving cells. To this end we expressed optoDop1R1^V2^ in the endogenous pattern of Dop1R1 using a knock-in Gal4 line (*Dop1R1^KO-Gal4^*). Locomotion of rotenone-treated larvae was tracked in the dark and subsequently upon green light illumination (Fig. 4B,C,movie S8). Strikingly, we observed light-dependent recovery of locomotion strictly dependent on optoDop1R1^V2^ function, which required feeding with 9-*cis*-Retinal. Optogenetic activation of Dop1R1 signaling significantly improved larval velocity and reduced overall turning behavior. This suggests that optoDop1R1^V2^ signaling in DA-receiving neurons can rescue toxin- induced dopaminergic impairment and corresponding locomotion deficits.

**Figure 4:**
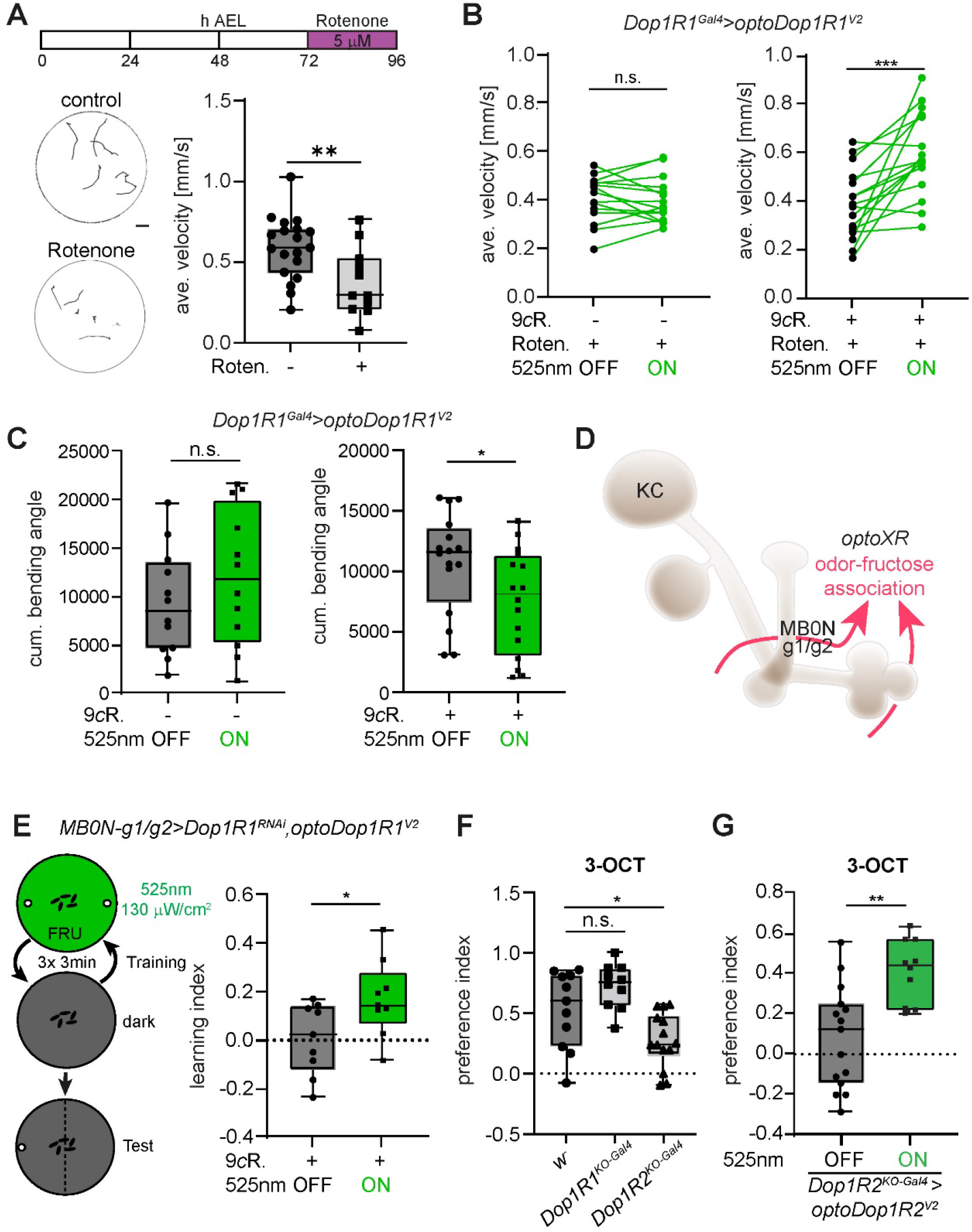
Functional validation of optoDopRs in *Drosophila* larvae *in vivo*. (A) Larvae were fed with Rotenone for 24h at 72 hours after egg laying (AEL) inducing locomotion defects due to impaired dopaminergic neuron function. Representative larval tracks of control or Rotenone-fed animals are shown (1 min, scale bar = 10 mm). Quantification of average velocity of control or Rotenone fed animals (n=19, 11, ** p<0.01). (B) Average velocity of Rotenone-fed animals expressing optoDop1R1^V2^ in an endogenous Dop1R1 pattern (*Dop1R1^ko-Gal4^>optoDop1R1^V^*^2^). Larvae were additionally fed with 9-*cis*-Retinal (9cR) and first tracked without light for 1min and with 525 nm light illumination for 1 min. Average velocity in the dark and during light activation is shown (n=15, 16, ** p<0.01). (C) Cumulative bending angles of Rotenone-fed animals expressing optoDop1R1^V2^ in an endogenous Dop1R1 pattern (*Dop1R1-Gal4>optoDop1R1^V^*^2^). Larvae without and with additional 9*-cis*- Retinal (9cR) feeding were tracked without light for 1min and with 525 nm light illumination for 1min. Bending angles in the dark and during light activation are shown (n=12, 16, * p<0.05). (D) Schematic model of the larval mushroom body consisting of Kenyon cells (KC) receiving input from dopaminergic neurons and connecting to output neurons (MBONs). Odor-Fructose association and learning requires dopaminergic input and MBON-g1/g2 (adapted from ^50^. (E) MBON-g1/g2 and Dop1R1-dependent single odor-fructose learning in larvae. Animals expressing optoDop1R1^V2^ and Dop1R1^RNAi^ in MBON- g1/g2 were trained using fructose-odor learning (3x3min) with or without light activation during fructose exposure (3 min 525 nm, 130 μW/cm^2^). Learning index of 9cR-fed animals with and without light activation during training are shown (n=9, 9,* p<0.05). (F) Innate preference for 3-Octanol (3-OCT) in control (w^-^), *Dop1R1^KO-Gal4^* and *Dop1R2^KO-Gal4^* 3^rd^ instar larvae (n=11, 10, 14, ***p<0.001). (G) Innate preference for 3-OCT in *Dop1R2^KO-Gal4^* 3^rd^ instar larvae expressing optoDop1R1^V2^. Innate preference for 3-OCT in 9cR-fed 3^rd^ instar animals with and without light activation during the assay (n=15, 10, * p<0.01).

We explored another core function of DA signaling by addressing its function in learning and memory. *Drosophila* larvae are capable of reward learning, e.g., by forming olfactory preferences through odor-fructose association ^38, 47^. As in adult flies, the MB plays a key role in this process: KCs receive specific DAergic input and form a tripartite circuit with MB output neurons (MBONs), which together reinforce specific preference behavior ^50^. Dop1R1 signaling and cAMP increase are essential for learning in flies. We therefore tested if optoDop1R1 activation during odor-fructose association can replace endogenous Dop1R1 function in KCs. We confirmed that KC-specific knockdown of Dop1R1 reduced learning performance in larvae (Fig. S4A). Using optoDop1R1^V1^ or optoDop1R1^V2^ expression in KCs under these conditions partially rescued fructose-odor learning (Fig. S4B,C). These results are consistent with the reported function of Dop1R1 in learning and suggest that acute activation of optoDop1R1 signaling in KCs during odor fructose-association is sufficient for learning. However, as dopaminergic responses in KCs were shown to be compartmentalized ^40, 51^, activation of optoDopRs in KCs cannot mimic this aspect of endogenous DA signaling. To avoid this issue, we tested for a potential function of Dop1R1 in MBON-g1/g2, which is specifically required for odor-fructose reward learning ^50^. RNAi-mediated knockdown of Dop1R1 in MBON-g1/g2 indeed reduced larval reward learning strongly suggesting DA signaling via Dop1R1 has an essential modulatory function in MBONs as well (Fig. S4D,E). We additionally expressed optoDop1R1^V2^ and activated it specifically during fructose-odor training, which partially rescued preference induction and learning compared to no light conditions (Fig. 4E, S4F). This suggests that acute optoDop1R1^V2^ activation during learning can functionally replace endogenous DA signaling in an MBON essential for odor-fructose association.

We further tested if DopRs are required for innate odor preference of Amylacetate (AM) and 3-Octanol (3-OCT), substances commonly used for larval odor-reward learning ^68, 69^. Dop1R1 knockout (*Dop1R1^ko-Gal4^*) and Dop1R2 knockout (*Dop1R2^ko-Gal4^*) larvae displayed no altered preference towards AM, which we used in our odor-reward learning paradigm (Fig. S4G).

However, *Dop1R2^ko^* larvae showed a specific reduction in 3-OCT preference (Fig. 4F). We therefore tested if optoDop1R2^V2^ activation could rescue innate preference behavior. Light exposure during the preference assay indeed was able to restore 3-OCT preference in *Dop1R2^ko-Gal4^* larvae expressing optoDop1R2^V2^ in an endogenous-like pattern (Fig. 4G). This result confirmed the functionality of optoDop1R2^V2^ by restoring the *in vivo* function of its corresponding wildtype receptor in odor preference.

### Functional analysis of dopaminergic signaling in adult flies

We further investigated the functionality of optoDopRs in adult flies, which requires very high light sensitivity of the optogenetic tools due to the low light penetrance of the fly cuticle, particularly below a wavelength of 530 nm ^70^. We first tested optoDop1R1^V2^ function in the MB in an associative odor-shock learning paradigm, which requires dopaminergic input from PPL1 neurons to KCs ^33, 71^. We confirmed that Dop1R1 is required in KCs for odor-shock learning using a MB-specific RNAi-mediated knockdown (Fig. 5A,B). We then asked if activation of optoDop1R1^V2^ in KCs can enhance performance when paired with the shock paradigm. We observed a trend towards more robust learning when optoDop1R1^V2^ was activated during shock pairing, but this performance was not significantly enhanced (Fig. 5A,C). Interestingly, optoDop1R1 co-activation reduced trial-dependent variability in this assay indicating more robust learning. We then asked if activation of DA signaling in KCs via optoDop1R1^V2^ activation could replace the shock stimulus, which would imply that this artificial DA signaling could replace a teaching signal with a negative valence. However, optogenetic activation of DA signaling without the unconditioned stimulus did not confer any preference behavior. These results indicate that either activation of Dop1R1 signaling alone is not sufficient for associative preference behavior or that the missing restriction to a distinct KC compartment interferes with memory formation.

**Figure 5:**
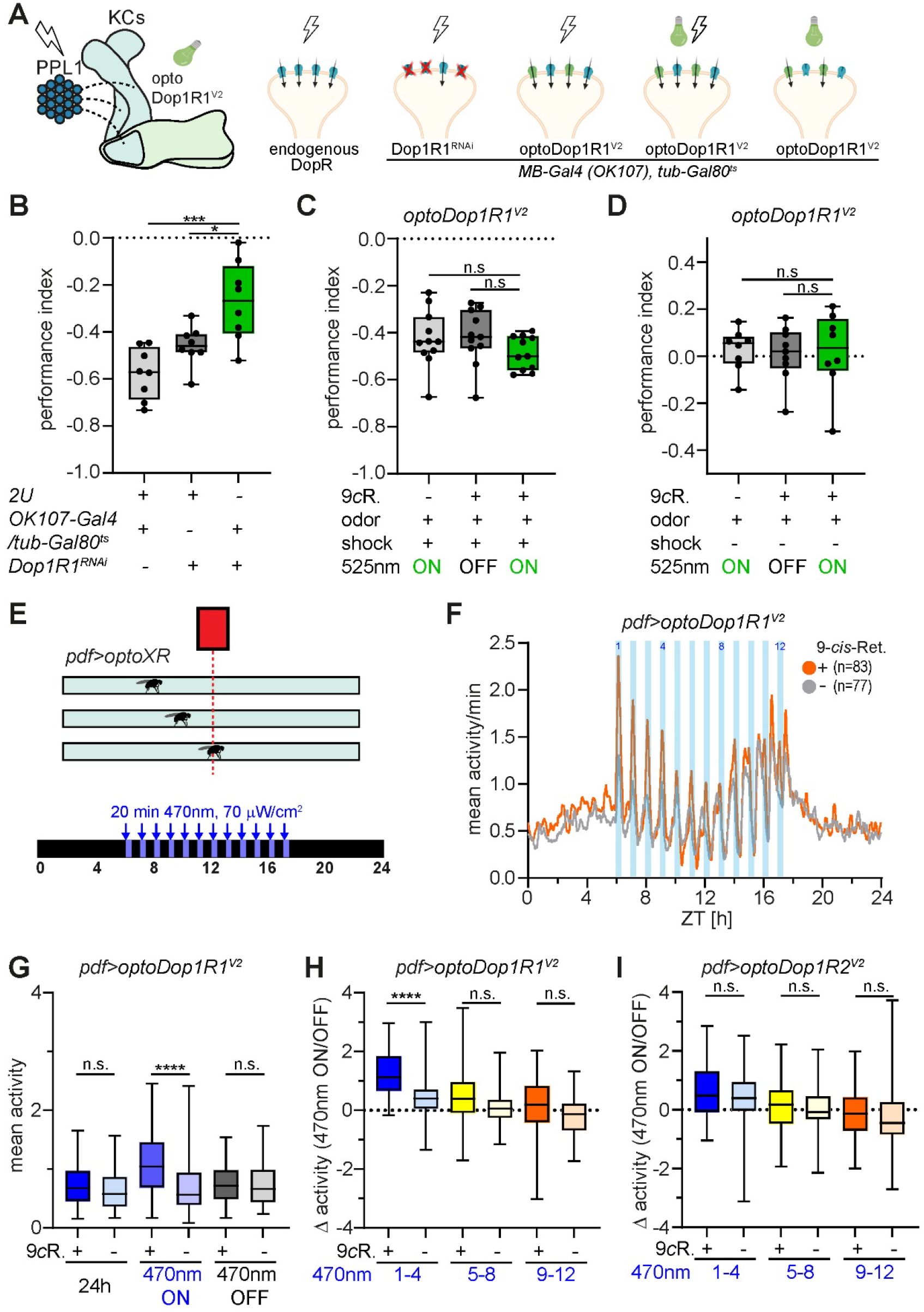
Cell type-specific function of acute Dop1R1 activity in adult flies. (A) Schematic of adult mushroom body organization and associative odor-shock learning under different conditions (shock and/or light and altered Dop1R1 activity). Adult flies (*OK107-Gal4; tub- Gal80^ts^; UAS-RNAi/optoXR*) for experiments in B-D were shifted to the permissive temperature (31°C) 4 days prior to the behavioral assay to induce Gal4 expression. (B) Performance index after aversive odor-shock learning with or without adult-specific RNAi-mediated knockdown of Dop1R1 in Kenyon cells (n=8, p<0.05, p<0.001). (C) Performance index after aversive odor-shock learning with or without additional activation of optoDop1R1^V2^ in Kenyon cells (n=11, p>0.05). (D) Performance index after pairing odor and optoDop1R1^V2^ activation in Kenyon cells (n=8, p>0.05). optoDop1R1^V2^-induced signaling is not sufficient to induce learning without the unconditioned stimulus. (E) Schematic of activity monitor with flies expressing optoXRs in pdf neurons with daytime-dependent light activation using a blue light stimulus. The grey bar indicates the flies’ subjective day. (F) Mean activity during 24h monitoring in flies expressing optoDop1R1^V2^ in pdf neurons with and without 9cR feeding (n: number of tested animals). Blue light pulses (12x 20min, 1/h) during subjective daytime increase fly activity during the morning hours. (G) Mean activity of *pdf>optoDop1R1^V^*^2^ -expressing flies during the entire 24h, all light on and light off phases (n=83,77, p<0.0001). (H) Activity difference of flies expressing optoDop1R1^V2^ in pdf neurons (with and without 9cR feeding) during light on/off times in the morning (1-4), midday (5-8) and afternoon (9-12) (n=83,77, p<0.0001). (I) Activity difference of flies expressing optoDop1R2^V2^ in pdf neurons (with and without 9cR feeding) during light on/off times in the morning (1-4), midday (5-8) and afternoon (9-12) (n=90, p>0.05). Note that only optoDop1R1^V2^ but not optoDop1R2^V2^ activation in pdf neurons during morning hours boosts fly activity.

We then assayed DopR function in I-LN_v_ neurons, which are important for circadian clock function ^72, 73^. Previous studies suggested that Dop1R1 has a depolarizing function in I-LN_v_s, affecting sleep and arousal state (^74^). We assayed the activity of flies using the *Drosophila* Activity Monitor (DAM) system^75^ from TriKinetics (Fig. 5E). Young flies were transferred to constant darkness after they had been reared under a 12 h dark/12 h light cycle. On the third day, darkness was interrupted by 12 arousing blue light pulses of different duration (10min, 15min, 20min) given every hour for a period of 12h that was in phase with the previous light period. These blue light-pulses not only aroused the flies but additionally activated optoDopRs expressed in I-LN_v_ neurons. Interestingly, expression and activation of optoDop1R1^V2^ was able to boost activity during the blue light periods compared to isogenic controls not fed with 9-*cis*-Retinal (Fig. 5F,G). We performed a more detailed analysis as the activity peaks were increasingly desynchronized with the blue light pulses (occurring after the light pulses) during the second part of the day. This revealed a significant effect of optoDop1R1^V2^ activation specifically during the first 4h window (Fig. 5H). Next, we also tested optoDop1R2^V2^ activation under the same conditions but did not observe a significant effect on blue-light induced activity (Fig. 5I, S5A-C). These findings suggest a specific role for Dop1R1 signaling in I-LN_v_s promoting morning activity upon arousal.

Finally, we also addressed a potential function of DopRs in adult MBONs previously implicated in encoding behavioral valence in MB-dependent tasks ^46, 76^. We chose an optoPAD setup which allows operant optogenetic stimulation of flies during feeding using a closed-loop system ^77^. We expressed optoDopRs in relevant MBONs providing output of the γ5/β’2-compartments of the MB and activated DA signaling with green light pulses every time the flies were sipping food (Fig. 6A). Operant activation of optoDop1R2^V2^ resulted in a decreased sipping rate over time suggesting that Dop1R2 signaling reduced the feeding drive and/or preference for the offered food (Fig. 6B,C). In contrast, operant optoDop1R1^V2^ activation during feeding did not result in changed feeding behavior (Fig. 6D, S6). We further asked if the endogenous DopRs played a role in feeding in valence encoding MBONs. RNAi-mediated knockdown of Dop1R2 but not Dop1R1 in MBON-γ5/β’2 resulted in an increased feeding rate (Fig. 6E,F, S6B,C) suggesting a specific function for Dop1R2 in these MBONs in feeding-related behavior. Controls without expression of optoDopRs did not show altered feeding with or without operant light exposure (Fig. S6D-G).

**Figure 6:**
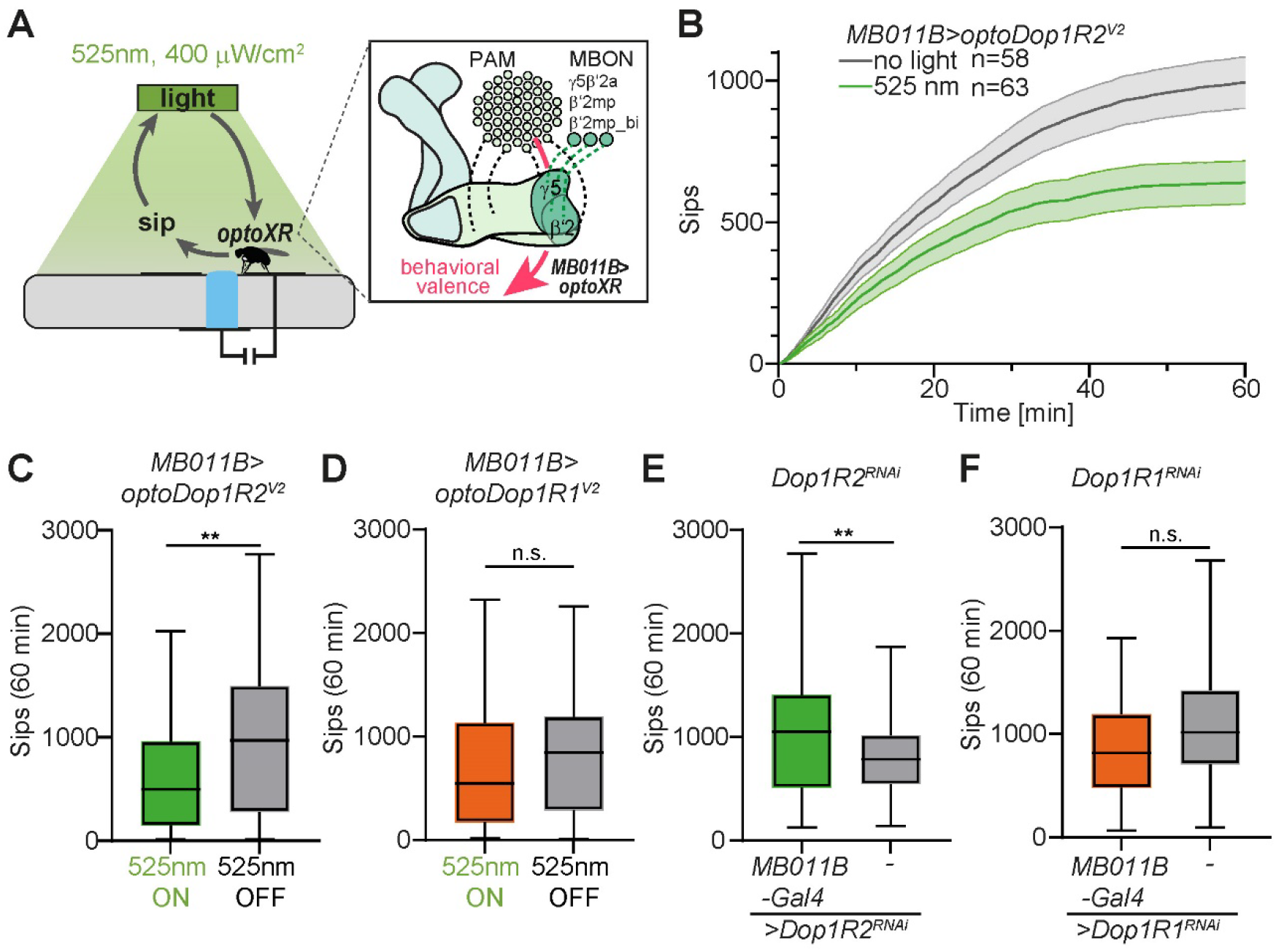
Cell type-specific function of operant Dop1R2 activity in adult satiety. (A) OptoPAD setup allowing light stimulation upon feeding action. Flies expressing optoXRs in a subset of MBONs (MBONγ5β’2a,β2mp and β2mp-bilateral) related to behavioral valence receive a light stimulus (1s, 525 nm 400 μW/cm^2^) every time they feed on the sucrose drop. (B) Cumulative sips over time for flies expressing optoDop1R2^V2^ using *MB011B-Gal4* with or without light stimulation (n=58,63). (C) Average sips at 60 min for flies expressing optoDop1R2^V2^ using *MB011B-Gal4* with or without light stimulation (n=58,63). (D) Average sips at 60min for flies expressing optoDop1R1^V2^ using *MB011B-Gal4* with or without light stimulation (n=65,65). (E) Average sips at 60min for flies expressing Dop1R2^RNAi^ with *MB011B-Gal4* and control (n=50,54). (F) Average sips at 60min for flies expressing Dop1R1^RNAi^ with *MB011B-Gal4* (n=47,41).

Taken together, operant optogenetic activation and RNAi-mediated decrease of Dop1R2 signaling in valence-encoding MBONs resulted in specific opposite effects on feeding. In contrast, manipulation of Dop1R1 activity in these MBONs did not alter feeding behavior. These findings strongly suggest that DA signaling in valence-encoding MBONs regulates feeding drive specifically via Dop1R2. Overall, these data show neuron-specific functions of Dop1R1 and Dop1R2 signaling which can be specifically induced by optoDopR activation.

## Discussion

By optimizing the chimeric optoXR approach we generated highly functional and specific optoDopRs that allowed *in vivo* analysis of receptor-specific function and behavior in *Drosophila*. optoDop1R1^V2^ showed enhanced and efficient activation in the blue and green light range up to 595nm in cellular assays with light-dose-dependent activation properties resembling the wildtype receptor. While Rho-based optoXRs display a broad spectral range of activation, they are compatible with red-shifted optogenetic tools including channelrhodopsins like Chrimson that can be activated above 600 nm ^56^. This should enable simultaneous optical control of neuronal activity *via* ion channel mediated as well as neuromodulatory pathways, providing a way forward towards all-optical access of neuronal network function *in vivo*. The high light sensitivity of the Rho backbone enables the activation of our optoXRs with blue or green light in adult flies *in vivo* despite less than 6% light penetrance of the adult cuticle in this spectral range ^70^. Although Rho is known to inactivate after its light cycle and only slowly being recycled ^78^, we did not observe a run-down in functionality *in vitro* or *in vivo*, possibly due to the abundance of the expressed optoXRs and the supplemented 9-*cis*-retinal.

Localization, cell-type specific and subcellular signaling dynamics are key to understanding endogenous GPCR signaling ^24, 79, 80^. Recent evidence showed that 2^nd^ messenger signaling occurs in nanodomains with receptor-specific profiles ^81^, emphasizing the importance of proper subcellular localization. Our optoDopR^V2^s display localization in the fly mushroom body in somatic and axonal compartments similar to their endogenous counterparts ^53^. In contrast, the previous design did not yield a functional optoDop1R2 receptor, and an optoDop1R1 mostly localizing to the somatic compartment with a signaling profile different from the wildtype receptor. This suggests that careful chimeric design is necessary to mimic endogenous receptor localization and function. This notion is consistent with the signaling profiles of both optoDopR^V2^s display specific signaling properties in KC axons mirroring the functions of their corresponding wildtype receptors. Dop1R1 has been shown to be required for cAMP responses in KCs, while Dop1R2 is required for calcium store release during olfactory conditioning ^37^. Therefore, these tools will be beneficial to further unravel their temporal activation requirements to induce functional associations during learning or goal-directed behavior.

DA signaling plays a complex role in innate and adaptive behaviors. We used a wide range of behavioral paradigms showing that our optoDopRs exhibit cell type, receptor, and behavioral paradigm-specific functions *in vivo*. We showed that both optoDopR^V2^s are functional and can at least partially replace endogenous DopRs in several assays including odor preference, locomotion and learning. At the same time, we uncovered a cell type specific requirement of DopR signaling: only optoDop1R1^V2^ but not optoDop1R2^V2^ activation promoted LN_v_-mediated arousal; *vice versa*, operant activation of optoDop1R2^V^^2^ but not optoDop1R1^V2^ in valence-encoding MBONs was able to control feeding. DopR function has been extensively studied in KCs but has so far not been investigated in MBONs. Our findings therefore strongly suggest that corresponding MB outputs are also under the control of DA signaling. Thus, our optoXRs provide an entry point to gain insight into temporal and cell type-specific DA signaling requirements of the insect learning center, enabling detailed studies of the temporospatial requirement of DA signaling for learning, valence encoding, goal-directed and innate behavior in one of the most developed and heavily used model systems.

Although our improved optoXR design allowed the generation of optoDopRs that are functional *in vivo*, the complexity of GPCR signaling and high sequence diversity of class A receptors makes general rational design of such tools difficult. Our incorporated adjustments provide an improved starting point that could be useful to generate optoXRs from other target receptors. Recently used approaches using structure-based design allowed improving the functionality of optoβ2AR, significantly increasing its light induced signaling properties ^17^.

However, experimental structures of optoDopRs are currently not available. Similarly, the implementation of spectrally tuned or bistable rhodopsin backbones, as for example shown for mouse Opn4 ^20, 82^, lamprey parapinopsin or mosquito Opn3 ^83^, yield further promise to extend the optoXR toolbox. Combinations of these complementary methods could further improve optoXR design and functionality to enable efficient chimera generation allowing *in vivo* studies of other receptors in the future.

## Acknowledgements

We thank A. Thum for *H24-Gal4* and *MBONg1/g2-Gal4*, M. Schwärzel for *UAS-bPAC*, J. Chu and Y. Li for *UAS-Gflamp1* fly lines. A. Schoofs for technical help with 2-Photon imaging. cDNA was obtained from the Drosophila Genomics Resource Center, supported by NIH grant 2P40OD010949. Stocks obtained from the Bloomington Drosophila Stock Center (NIH P40OD018537) were used in this study.

## Author Contributions

FZ performed in vitro characterization and analysis of DopRs/optoDopRs, cAMP signaling and larval behavior, AMT and HJ designed optoDopRs, BNI performed and analyzed functional in vivo imaging, FJRJ, MGM,IJ and IGK designed, performed and analyzed flyPAD/optoPAD experiments, MH and CHF designed, performed and analyzed locomotor activity experiments, NW, VB and JP designed, performed and analyzed adult learning experiments, TL and KS contributed to in vitro characterization and analysis of DopRs/optoDopRs, PS designed the study and wrote the manuscript with input from all authors.

## Materials & Correspondence

Correspondence and material requests should be addressed to

## Methods

### OptoDopR design

OptoDopR sequences were designed using bovine rhodopsin as the acceptor receptor with G-protein binding sites exchanged for those of the target receptor. To determine cut sites, a multiple protein sequence alignment of bovine rhodopsin and the target receptors was generated using muscle^84^. Macros written in IgorPro were then used to cut and combine the aligned protein sequences in an automated fashion. V1 cut sites were based on previously published receptor designs^14, 18^. For V2, cut sites around ICL1 and the C-terminus were amended to reflect previously published insights on G-protein binding sites^43^: residues in ICL1 were shown to not contribute to G-protein binding, thus cut sites in ICL1 were removed to retain rhodopsin ICL1. Correspondingly, the C-terminal cut sites were moved further towards the TM domains as these residues were shown to contribute to G-protein binding.

### Plasmids

cDNAs of wildtype Drosophila Dop1R1 and Dop1R2 were obtained from the Drosophila Genomics Resource Center (DGRC, Bloomington, IN, USA) and cloned into pCDNA3.1 (ThermoFisher, MA, USA). optoDop1R1 and optoDop1R2 chimera (V1 and V2) were synthesized as codon-optimized cDNAs (ThermoFischer, MA, USA) and cloned into pCDNA3.1 and pUAttB. Chimeric G proteins for the G_sx_ assay ^44^ were obtained from Addgene (Watertown, MA, USA).

### Cell culture and live-cell cAMP assay

G protein coupling of wild type and chimeric GPCR constructs was tested in HEK293T cells (or HEK293T-dd7 cells) by using a live cell assay ^44^. The GPCR constructs were subcloned into pcDNA3.1 (ThermoFisher, MA, USA). HEK293T cells were incubated in DMEM medium containing 10% FBS (FAN Tech.) with penicillin (100 U/mL) and streptomycin (100 mg/mL) at 37°C and 5% CO_2_. For transfection, cells were seeded into white 96-well plates (Greiner Bio One) coated with poly-L-lysine (Sigma Aldrich, St. Louis, MO, USA) and incubated at 37°C for 1 hour before being transfected with individual receptors, G protein chimera (G_sx_) and Glo22F (Promega) using Lipofectamine 2000 (ThermoFisher, MA, USA).

Cells were incubated at 37°C and 5% CO_2_ for 24 h and the medium replaced with L-15 media (without phenol-red, 1% FBS) containing 2 mM beetle luciferin (in 10 mM HEPES pH 6.9) and 10 mM 9-*cis*-retinal (for optoXRs) and cells were incubated at room temperature for 1h. For optoXR experiments, the plates were kept in the dark at all times before illumination and cAMP dependent luminescence was measured using a Berthold Mithras multimode plate reader (Berthold Tech., Germany). Baseline luminescence was measured three times and activation of DopRs was induced by ligand addition (dopamine at various concentrations diluted in L-15). For optoDopR activation cells were illuminated with a 1 second light pulse using an LED light plate (Phlox Corp., Provence, France) or a CoolLED pE-4000 (CoolLED, Andover, UK). Specific light intensities and wavelengths are indicated in individual experiments. Technical duplicates were performed for all experiments with a minimum of three independent trials. For data quantification each well was normalized to its pre-activation baseline.

### Drosophila melanogaster stocks

All *Drosophila* stocks were raised and treated under standard conditions at 25 °C and 70% relative humidity with a 12 h light/dark cycle on standard fly food unless stated otherwise. Transgenic UAS-optoDopR lines were generated by phiC31-mediated site-specific transgene using the attP2 site on the 3rd chromosome (FlyORF Injection Service, Zurich, Switzerland). Stocks were obtained from the Bloomington (BDSC) Drosophila stock centers, unless otherwise noted. We used the lines as shown in the Table 1.

**Table 1.** Lines used in this study

| Line | label | Source |
| --- | --- | --- |
| *Dop1R1<sup>KO-Gal4</sup>* | Knockout-Gal4 of Dop1R1 | BDSC# 84714 |
| *UAS-Dop1R1<sup>RNAi</sup>* | Knockdown of Dop1R1 | BDSC# 62193 |
| *UAS-Dop1R2<sup>RNAi</sup>* | Knockdown of Dop1R2 | BDSC# 51423 |
| *Dop1R2<sup>KO-Gal4</sup>* | Knockout-Gal4 of Dop1R2 | BDSC# 84715 |
| *201y-Gal4* | Expresses GAL4 in the mushroom body | BDSC# 64296 |
| *H24-Gal4* | Expresses GAL4 in the mushroom body | BDSC# 51632 |
| *UAS-bPAC* | Optogenetic activation | <sup>55</sup>, BDSC# 78788 |
| *UAS-Dop1R1<sup>V2</sup>* | Optogenetic activation | This study |
| *UAS-Dop1R2<sup>V2</sup>* | Optogenetic activation | This study |
| *UAS-Dop1R1<sup>V1</sup>* | Optogenetic activation | This study |
| *ppk-Gal4* | Expresses GAL4 in C4da neurons | <sup>85</sup> |
| *UAS-CsChrimson-GFP* | Optogenetic activation | <sup>56</sup>, BDSC# 55136 |
| *UAS-Gflamp1* | cAMP reporter | <sup>57</sup> |
| *UAS-Gcamp6s* | calcium reporter | <sup>59</sup> |
| *MBONg1g2-Gal4* | Expresses GAL4 in MBON-g1,g2 | <sup>50</sup> |
| *Pdf-Gal4* | Expresses GAL4 in I-LN<sub>v</sub> | BDSC# 6899 |
| *MB011B-Gal4* | Expresses GAL4 in valence-encoding MBONs | <sup>76</sup> |
| *2U* | w<sup>1118</sup> (isoCJ1) Canton-S derivative | <sup>86</sup> |
| *OK107-Gal4* | Expresses GAL4 in the mushroom body | BDSC# 854 |
| *tub-Gal80<sup>ts</sup>* | Expresses temperature sensitive GAL80 in all cells | BDSC# 7019 |
| *R21B06-splitGal4<sup>DBD</sup>* | Expresses GAL4<sup>DBD</sup> in the mushroom body | <sup>46</sup> |
| *6xCRE-splitGal4<sup>AD</sup>* | Expresses GAL4<sup>AD</sup> in a Cre-dependent manner, VK27 insertion | This study, see <sup>52</sup> |
| *UAS-myr::tdTomato* | Fluorescent reporter line | <sup>87</sup>, BDSC# 32223 |

### Immunochemistry

Larval brains from 3^rd^ instar animals (96 h ±3h AEL) of the indicated genotypes were dissected in phosphate-buffered saline (PBS) and fixed for 15min at room temperature in 4% paraformaldehyde/PBS. After washing in PBST (PBS with 0.3% Triton X-100) and incubation in 5% normal donkey serum in PBST, the samples were incubated with mouse anti-Rhodopsin antibody (1:100) at 4°C overnight, washed in PBST 3 times (5 min in each time) and incubated with secondary antibodies (mouse anti-Cy3, Jackson Immunoresearch,1:300) for 1h. After washing, samples were mounted on poly-L-lysine coated coverslips in Slow Fade Gold (Thermo Fisher, CA, USA). Native reporter fluorescence was sufficiently bright to be visualized together with antibody immunostaining by confocal microscopy (Zeiss LSM900AS2, Zeiss, Oberkochen, Germany). Confocal Z stacks were processed in Fiji (ImageJ, NIH, Bethesda, USA).

Adult brains of 3-7 days old flies of the indicated genotypes were dissected in hemolymph-like saline (HL3) and fixed for 1 h at room temperature in 2 % paraformaldehyde/HL3. After washing in PBST (PBS with 0.5 % Triton X-100) and incubation in 5 % normal goat serum in PBST, samples were incubated with mouse anti-Rhodopsin (1D4, 1:1,000, Thermo Fisher), rabbit anti-DsRed (1:2,000, Takara Bio Inc.), rabbit or guinea pig anti-Discs large (1:30,000 and 1:1.000; ^88^) antibodies for 4 h at room temperature, followed by 2 nights at 4°C. Samples were subsequently washed in PBST (3 x 30 min) and incubated with secondary antibodies (goat anti-mouse Alexa488, goat anti-rabbit Alexa 594, goat anti-guinea pig Alexa 647, 1:1,000, Thermo Fisher) for 4 h at room temperature, followed by 2 nights at 4°C. After washing, a pre-embedding fixation in 4 % paraformaldehyde/PBS was performed for 4 h at room temperature. Samples were washed in PBST (4 x 15 min) followed by 10 min in PBS. Brains were mounted on poly-L-lysine coated coverslips. An ethanol dehydration series and a xylene clearing series were performed and the samples were mounted in DPX ^89^. Images were taken on a Leica STELLARIS 8 confocal microscope using a 20x (NA 0.75) and 93x (NA 1.3) glycerol immersion objective. Confocal z-stacks were processed in Fiji (ImageJ, NIH, Bethesda, USA).

### Calcium and cAMP imaging in D. melanogaster larvae

3^rd^ instar larval brains (96 h ±3h AEL) were partially dissected in physiological saline buffer (108 mM NaCl, 5mM KCl, 2mM CaCl_2_, 8.2 mM MgCl_2_, 4 mM NaHCO_3_, 1 mM NaH_2_PO_4_, 5 mM trehalose, 10 mM sucrose, 5 mM HEPES, pH 7.5) and mounted on poly-L-lysine -coated cover slips in the saline buffer with or without 5mM 9-*cis*-Retinal (for opto-Dop1R1). Gflamp-1 was utilized to monitor cAMP levels *in vivo*. Live imaging of Kenyon cell somata and medial lobes expressing Gflamp-1 in the mushroom body was performed using confocal microscopy with a 40x/NA1.3 objective (Zeiss LSM900AS2, Zeiss, Oberkochen, Germany).

OptoDop1R1^V2^ or bPAC activation was achieved using 470 nm LED light with an intensity of 2.10 mW/cm². Confocal time series were recorded at 7.5 frames/s (128 × 128 pixels, 600 frames total). KC somata or medial lobes were focused and after a stable imaging period of 100 frames the 470 nm LED was activated for 10 seconds. Confocal time series were analyzed using image registration (StackReg plugin, ImageJ) to correct for XY movement and Gflamp-1 signal intensity in the soma and medium lobe was quantified using the Time Series Analyzer V3 plugin (ImageJ). Baseline (F_0_) was determined as the average of 95 frames before activation. The relative maximum intensity change (ΔF_max_) of Gflamp-1 fluorescence was calculated after normalization to baseline.

Calcium responses were recorded from the soma/calyx region and the medial lobe of the mushroom body using *UAS-GCaMP6s* and *UAS-OptoDop1R2* ^V2^ under the control of H*24-Gal4*. Animals were reared in the dark on grape agar plates supplemented with yeast paste and 9-*cis*-retinal. Live imaging on 3^rd^ instar larvae was performed under low light conditions. Larvae were mounted in 90% glycerol, sandwiched between a coverslip and the slide with the aid of silicon paste. The soma as well as the medial lobe of the mushroom body were live imaged using a Zeiss LSM 780 2-photon microscope and a 25x/NA1.0 water immersion objective. For activation of the optoDop1R2^V2^, larvae were subjected to 10s blue light stimulation (470 nm, 720 μW/cm², CoolLED) twice with an interval of 30s between each pulse. Only datasets without significant Z-drift were used for analysis. Analysis of the time series was performed using Fiji (ImageJ, NIH, Bethesda, USA) as described above.

Normalized calcium responses were obtained by subtracting the amplitude of the pre-stimulation baseline (average of 50 frames) from the stimulation evoked amplitude. The calcium response was recorded before and after the light stimulus due to PMT overexposure during the light pulse. Graphs showing the mean ± s.e.m were generated with GraphPad Prism (GraphPad, San Diego, CA, USA). Boxplots was used to show the comparison between the maximum responses (ΔF_max_/F_0_) and analyzed with unpaired Student’s t-test with Welch’s correction.

### cAMP-induced nociceptive behavior in D. melanogaster larvae

For cAMP-induced nociceptive behavior, larvae expressing UAS-bPAC, UAS-CsChrimson or UAS-optoDopRs under the control of *ppk-Gal4* were staged and fed in the dark on grape agar plates (2% agar) with yeast paste containing 5 mM 9-*cis*-retinal (optoXRs) or all-*trans*- retinal (CsChrimson). Staged 3^rd^ instar larvae were placed on a 1% agar film on a FTIR (frustrated total internal reflection) based tracking system (FIM, University of Münster) with 1ml water added. Experiments were performed under minimum light conditions (no activation). After 10s, larvae were illuminated with 470 nm light (465 μW/cm²) for 3min.

Behavioral responses (rolling) were recorded, analyzed, and plotted. Staging and experiments were performed in a blinded and randomized manner.

### Locomotion assays in D. melanogaster larvae

*D. melanogaster* larvae were staged in darkness on grape agar plates containing yeast paste with or without 5 mM 9-*cis*-retinal. For the indicated experiments, larvae were additionally fed with Rotenone for 24h at 72 hours after egg laying (AEL) to impair dopaminergic neuron function. Third instar larvae (96 h ± 4 h AEL) were used for all experiments. Animals were carefully selected and transferred under minimum red-light conditions to a 1% agar film on a FTIR (frustrated total internal reflection) based tracking system (FIM, University of Münster). Five freely moving larvae per trial were video-captured and stimulated with 525 nm light (130 μW/cm²) for activation of optoDop1R1^V2^. Animal locomotion was tracked with 10 frames/s for up to 120s. For locomotion analysis, velocity and bending angles were analyzed using the FIMtrack software (https://github.com/kostasl/ FIMTrack). Only animals displaying continuous locomotion before the light stimulus were analyzed. Average locomotion speed and cumulative bending angles were analyzed and plotted for the first 30 s under dark or light conditions.

### Innate odor preference and olfactory behavior assays in D. melanogaster larvae

Groups of 20 staged mid-3^rd^ instar larvae (96h±4h AEL) were placed in the middle of a 2% agar plate containing a container with 10 µl n-amylacetate (AM, diluted 1:50 in mineral oil; SAFC) or 3-Octanol (3-Oct, Sigma) on one side and a blank on the other side. For rescue experiments, assays were performed either in the dark or using light conditions (525 nm, 130 μW/cm²) during the preference behavior. Assays were video captured for 5 min under infrared light illumination to monitor larval distribution with a digital camera (Basler ace-2040 gm, Basler, Switzerland). After 5 min the number of larvae on each side was determined and the odor preference was calculated as (n(larvae) on odor side – n(larvae) on blank side)/total n(larvae).

### Odor-fructose reward learning assays in D. melanogaster larvae

Odor-fructose reward learning was performed essentially as described ^69^. Groups of 20 larvae each were placed in a petri dish coated either with plain 2% agar or 2% agar with 2 M fructose as a reward in the presence of 10 µl n-amylacetate (AM, 1:50). The odor-reward or no reward pairing was done for 3 min (or 5min; as indicated in experiments), alternating 3x between training (odor^+^), while the unpaired group received odor and reward during separate 3min (or 5min as indicated) training (blank^+^). For all optogenetic lines, training was performed under minimum red-light conditions, or with 525nm light activation (130 μW/cm²) during fructose reward training. Reciprocal training was performed for all genotypes and conditions (blank/odor^+^ and blank^+^/odor, respectively).

After three training cycles, larval preference towards the trained odor (AM or blank) was recorded in darkness using a Basler ace-2040gm camera (same setting as for the olfactory behavior assay). The number of larvae on each side was calculated after 5 minutes, and odor preferences were calculated for the paired and unpaired groups. The learning index (LI) was then calculated using the following formula:

LI = (Odor- Pref _Paired_ – Odor -Pref _Unpaired_)/2

### Odor-shock learning behavior assays in D. melanogaster adult flies

Aversive olfactory conditioning of adult flies was performed as described before ^69^. Conditioning was performed in the dark at 21°C and 75 % humidity using 3-7 day old flies. Groups of flies were loaded into custom-made copper grid tubes with high power LEDs mounted at the end of the tube (525 nm, Ø 37 µW/mm²). Flies were exposed to a constant air stream or the odorized air stream (750 ml/min).

Experimental flies were raised at 20 °C and shifted to 31 °C four days prior to the experiments to induce Gal80^ts^/ Gal4-dependent gene expression. Flies were transferred to 0.4 mM 9-*cis*-retinal food ∼ 48 h prior to the experiment and kept in the dark.

For conditioning the odors 4-MCH (1:250, Merck, Darmstadt, Germany,CAS #589-91-3) and 3-OCT (1:167, Merck, Darmstadt, Germany, CAS #589-98-0) were diluted in mineral oil (Thermo Fisher, Waltham, MA, CAS #8042-47-5). Flies were conditioned following a five times spaced training paradigm. After a resting period of 3 min with only airflow the flies were exposed to the stimuli as indicated in the figure. The CS^+^, electric shocks (twelve 1.5-second 90 V shocks with 3.5-second intervals) (Fig. 5B) and pulsed green light (4 Hz, 0.125 s on and 0.125 s off) (Fig. 5C, D) were applied simultaneously for 60 s. After 45 s of airflow the CS^-^ was presented for 60 s. This training cycle was repeated five times with 15 min breaks in between cycles. Odors for CS+ and CS- were interchanged for each n.

Learning behavior was subsequently analyzed in the T-Maze. At the decision point of the T-Maze flies could choose for 2 min between the CS^+^ and the CS^-^ (OCT 1:670, MCH 1:1,000). The performance index was calculated for MCH and OCT individually:

Performance index = (# of flies (CS^+^) - # of flies (CS^-^)) / total # of flies

For each n the two data points obtained with MCH and OCT as CS+ were averaged.

### DopR function in I-LNv neurons of D. melanogaster adults

Flies were raised under 12h:12h light-dark cycles at 20°C on standard fly food. 1-4 day old male flies were placed individually in DAM (TriKinetics) monitors ^75^ containing 2% agar with 4% sucrose and 5mM 9-*cis*-Retinal solved in ethanol (for opto-Dop1R1 and opto-Dop1R2) or only ethanol (for controls). The activity of the flies was recorded in complete darkness for 2 days before the flies were subjected to light-pulses of 470 nm LED light with an intensity of 70 ± 10 µW/cm². The light-pulses were administered 12 times during the previous light-period of the 12h:12h light-dark cycle (one pulse every hour for 10min, 15min or 20 min).

Experiments were performed 3 times with 32 experimental and control flies, respectively. Activity data were plotted as individual and average actograms using the ImageJ plug-in actogramJ ^90^, and individual and average activity profiles of the 24 h day with light pulses were calculated for each fly group as described in ^91^.

### Feeding behavior assays in D. melanogaster adults

Flies used in the flyPAD were reared and maintained in standard cornmeal food, with composition described before ^92^ in incubators at 28 °C, 60% humidity and cycles of light/dark of 12 hours each. After hatching, male flies of 4-8 days old were collected. 5 µl of 10% sucrose solution containing 1% low gelling temperature agarose were placed in wells of the flyPAD containing electrodes to detect the capacitance change when the flies physically interacted with the food. The flies, following starvation for 24 h in the presence of a wet tissue with 3 ml of water, were transferred to the flyPAD individually using a pump. The experiments were all performed in a climate chamber at 25 °C, at 60% humidity. The recording of each session of flyPAD lasted 60 minutes, in which the flies could freely interact with the food.

For the optoPAD experiments^77^, flies were reared and maintained in standard cornmeal food as explained above, with supplementation of all-trans-retinal at a 1:500 concentration, in incubators at 25 °C, 60% humidity and blue light/dark cycles of 12/12 hours. The chimeric dopaminergic receptors were activated using 523 nm green light, at 3 V, which was automatically activated once the fly started to sip food. All flies were wet starved for 24 h prior to the experiment. The acquisition of the data was done using scripts (https://github.com/ribeiro-lab/optoPAD-software) based on Bonsai, an open-source program.

The analysis was done using a Matlab code.

### Quantification and statistical analysis

Statistical analysis was performed using Prism 8 (Graphpad, San Diego, CA, USA). Boxplots depict the median with 25^th^ and 75^th^ percentile (lower and upper box, respectively) and whiskers represent the 1^st^ and 99^th^ percentile. For line graphs, the average and ±s.e.m. are shown. Unpaired two-tailed Student’s t-test (for two groups with normally distributed data), paired two-tailed Student’s t-test (for locomotion behavior rescue experiment), one-way ANOVA (for multiple comparisons), or χ² tests were used for group comparisons as indicated.

## References

1. Bargmann, C. I. & Marder, E. From the connectome to brain function. Nat Methods 10, 483–490 (2013).

2. Klein, M. O., et al. Dopamine: Functions, Signaling, and Association with Neurological Diseases. Cellular and Molecular Neurobiology 2018 39:1 39, 31–59 (2018).

3. Girault, J. A. & Greengard, P. The Neurobiology of Dopamine Signaling. Arch Neurol 61, 641–644 (2004).

4. Beaulieu, J. M. & Gainetdinov, R. R. The physiology, signaling, and pharmacology of dopamine receptors. Pharmacol Rev 63, 182–217 (2011).

5. Siju, K. P., De Backer, J. F. & Grunwald Kadow, I. C. Dopamine modulation of sensory processing and adaptive behavior in flies. Cell and Tissue Research 2021 383:1 383, 207–225 (2021).

6. Karam, C. S., Jones, S. K. & Javitch, J. A. Come Fly with Me: An overview of dopamine receptors in Drosophila melanogaster. Basic Clin Pharmacol Toxicol 1, 1– 10 (2019).

7. Deisseroth, K. Optogenetics: 10 years of microbial opsins in neuroscience. Nat Neurosci 18, 1213–1225 (2015).

8. Deisseroth, K. Optogenetics. Nat Methods 8, 26–29 (2011).

9. Wiegert, J. S., Mahn, M., Prigge, M., Printz, Y. & Yizhar, O. Silencing Neurons: Tools, Applications, and Experimental Constraints. Neuron 95, 504–529 (2017).

10. Tichy, A.-M., Gerrard, E. J., Sexton, P. M. & Janovjak, H. Light-activated chimeric GPCRs: limitations and opportunities. Curr Opin Struct Biol 57, 196–203 (2019).

11. Spangler, S. M. & Bruchas, M. R. Optogenetic approaches for dissecting neuromodulation and GPCR signaling in neural circuits. Curr Opin Pharmacol 32, 56– 70 (2017).

12. Rost, B. R., Schneider-Warme, F., Schmitz, D. & Hegemann, P. Optogenetic Tools for Subcellular Applications in Neuroscience. Neuron 96, 572–603 (2017).

13. Siuda, E. R. et al. Optodynamic simulation of β-adrenergic receptor signalling. Nat Commun 6, 8480 (2015).

14. Kim, J.-M. et al. Light-Driven Activation of β 2 -Adrenergic Receptor Signaling by a Chimeric Rhodopsin Containing the β 2 -Adrenergic Receptor Cytoplasmic Loops †. Biochemistry 44, 2284–2292 (2005).

15. Bailes, H. J., Zhuang, L.-Y. & Lucas, R. J. Reproducible and Sustained Regulation of Gαs Signalling Using a Metazoan Opsin as an Optogenetic Tool. PLoS One 7, e30774 (2012).

16. Airan, R. D., Thompson, K. R., Fenno, L. E., Bernstein, H. & Deisseroth, K. Temporally precise in vivo control of intracellular signalling. Nature 458, 1025–1029 (2009).

17. Tichy, A. M., So, W. L., Gerrard, E. J. & Janovjak, H. Structure-guided optimization of light-activated chimeric G-protein-coupled receptors. Structure 30, 1075–1087.e4 (2022).

18. Morri, M. et al. Optical functionalization of human Class A orphan G-protein-coupled receptors. Nat Commun 9, 1950 (2018).

19. Čapek, D. et al. Light-activated Frizzled7 reveals a permissive role of non-canonical wnt signaling in mesendoderm cell migration. Elife 8, (2019).

20. van Wyk, M., Pielecka-Fortuna, J., Löwel, S. & Kleinlogel, S. Restoring the ON Switch in Blind Retinas: Opto-mGluR6, a Next-Generation, Cell-Tailored Optogenetic Tool. PLoS Biol 13, 1–30 (2015).

21. Eichel, K. & von Zastrow, M. Subcellular Organization of GPCR Signaling. Trends Pharmacol Sci 39, 200–208 (2018).

22. Spoida, K., Masseck, O. A., Deneris, E. S. & Herlitze, S. Gq/5-HT2c receptor signals activate a local GABAergic inhibitory feedback circuit to modulate serotonergic firing and anxiety in mice. Proc Natl Acad Sci U S A 111, 6479–6484 (2014).

23. Eickelbeck, D. et al. CaMello-XR enables visualization and optogenetic control of Gq/11 signals and receptor trafficking in GPCR-specific domains. Communications Biology 2019 2:1 2, 1–16 (2019).

24. Lohse, M. J. & Hofmann, K. P. Spatial and Temporal Aspects of Signaling by G- Protein–Coupled Receptors. Mol Pharmacol 88, 572–578 (2015).

25. Nässel, D. R. Substrates for Neuronal Cotransmission With Neuropeptides and Small Molecule Neurotransmitters in Drosophila. Front Cell Neurosci 12, 1–26 (2018).

26. Nässel, D. R., Pauls, D. & Huetteroth, W. Neuropeptides in modulation of Drosophila behavior : how to get a grip on their pleiotropic actions. 1–19 (2019) doi:10.7287/peerj.preprints.27531v1.

27. Bargmann, C. I. Beyond the connectome: How neuromodulators shape neural circuits. BioEssays 34, 458–465 (2012).

28. Taghert, P. H. & Nitabach, M. N. Peptide neuromodulation in invertebrate model systems. Neuron 76, 82–97 (2012).

29. Nässel, D. R. & Zandawala, M. Endocrine cybernetics: neuropeptides as molecular switches in behavioural decisions. Open Biol 12, 24–26 (2022).

30. Zolin, A. et al. Context-dependent representations of movement in Drosophila dopaminergic reinforcement pathways. Nat Neurosci 24, 1555–1566 (2021).

31. Kaun, K. R. & Rothenfluh, A. Dopaminergic rules of engagement for memory in Drosophila. Curr Opin Neurobiol 43, 56–62 (2017).

32. Waddell, S. Reinforcement signalling in Drosophila; dopamine does it all after all. Curr Opin Neurobiol 23, 324–9 (2013).

33. Adel, M. & Griffith, L. C. The Role of Dopamine in Associative Learning in Drosophila: An Updated Unified Model. Neurosci Bull 37, 831–852 (2021).

34. Berry, J. A., Cervantes-Sandoval, I., Nicholas, E. P. & Davis, R. L. Dopamine is required for learning and forgetting in Drosophila. Neuron 74, 530–42 (2012).

35. Keleman, K. et al. Dopamine neurons modulate pheromone responses in Drosophila courtship learning. Nature 489, 145–9 (2012).

36. Burke, C. J. et al. Layered reward signalling through octopamine and dopamine in Drosophila. Nature 492, 433–7 (2012).

37. Handler, A. et al. Distinct Dopamine Receptor Pathways Underlie the Temporal Sensitivity of Associative Learning. Cell 178, 60–75.e19 (2019).

38. Rohwedder, A. et al. Four Individually Identified Paired Dopamine Neurons Signal Reward in Larval Drosophila. Current Biology 26, 661–669 (2016).

39. Himmelreich, S. et al. Dopamine Receptor DAMB Signals via Gq to Mediate Forgetting in Drosophila. Cell Rep 21, 2074–2081 (2017).

40. Boto, T., Louis, T., Jindachomthong, K., Jalink, K. & Tomchik, S. M. Dopaminergic Modulation of cAMP Drives Nonlinear Plasticity across the Drosophila Mushroom Body Lobes. Current Biology 24, 822–831 (2014).

41. Hickey, D. G. et al. Chimeric human opsins as optogenetic light sensitisers. J Exp Biol 224, (2021).

42. Gunaydin, L. A. et al. Natural Neural Projection Dynamics Underlying Social Behavior. Cell 157, 1535–1551 (2014).

43. Flock, T. et al. Selectivity determinants of GPCR-G-protein binding. Nature 545, 317– 322 (2017).

44. Ballister, E. R., Rodgers, J., Martial, F. & Lucas, R. J. A live cell assay of GPCR coupling allows identification of optogenetic tools for controlling Go and Gi signaling. BMC Biol 16, 10 (2018).

45. Heisenberg, M. Mushroom body memoir: from maps to models. Nat Rev Neurosci 4, 266–75 (2003).

46. Aso, Y. et al. The neuronal architecture of the mushroom body provides a logic for associative learning. Elife 3, e04577 (2014).

47. Thum, A. S. & Gerber, B. Connectomics and function of a memory network: the mushroom body of larval Drosophila. Curr Opin Neurobiol 54, 146–154 (2019).

48. Eichler, K. et al. The complete connectome of a learning and memory centre in an insect brain. Nature 548, 175–182 (2017).

49. Eschbach, C. et al. Recurrent architecture for adaptive regulation of learning in the insect brain. Nat Neurosci 23, 544–555 (2020).

50. Saumweber, T. et al. Functional architecture of reward learning in mushroom body extrinsic neurons of larval Drosophila. Nat Commun 9, 1104 (2018).

51. Cohn, R., Morantte, I. & Ruta, V. Coordinated and Compartmentalized Neuromodulation Shapes Sensory Processing in Drosophila. Cell 163, 1742–1755 (2015).

52. Siegenthaler, D., Escribano, B., Bräuler, V. & Pielage, J. Selective suppression and recall of long-term memories in Drosophila. PLoS Biol 17, e3000400 (2019).

53. Kondo, S. et al. Neurochemical Organization of the Drosophila Brain Visualized by Endogenously Tagged Neurotransmitter Receptors. Cell Rep 30, 284–297.e5 (2020).

54. Dannhäuser, S. et al. Antinociceptive modulation by the adhesion GPCR CIRL promotes mechanosensory signal discrimination. Elife 9, (2020).

55. Stierl, M. et al. Light modulation of cellular cAMP by a small bacterial photoactivated adenylyl cyclase, bPAC, of the soil bacterium Beggiatoa. J Biol Chem 286, 1181–8 (2011).

56. Klapoetke, N. C. et al. Independent optical excitation of distinct neural populations. Nat Methods 11, 338–46 (2014).

57. Wang, L. et al. A high-performance genetically encoded fluorescent indicator for in vivo cAMP imaging. Nat Commun 13, 5363 (2022).

58. Yang, S. et al. PACmn for improved optogenetic control of intracellular cAMP. BMC Biol 19, 1–17 (2021).

59. Chen, T.-W. et al. Ultrasensitive fluorescent proteins for imaging neuronal activity. Nature 499, 295–300 (2013).

60. Glovaci, I. & Chapman, C. A. Dopamine induces release of calcium from internal stores in layer II lateral entorhinal cortex fan cells. Cell Calcium 80, 103–111 (2019).

61. Sayin, S. et al. A Neural Circuit Arbitrates between Persistence and Withdrawal in Hungry Drosophila. Neuron 104, 544–558.e6 (2019).

62. Lewis, L. P. C. et al. A Higher Brain Circuit for Immediate Integration of Conflicting Sensory Information in Drosophila. Current Biology 25, 2203–2214 (2015).

63. Montgomery, E. B. Basal ganglia pathophysiology in Parkinson’s disease. Ann. Neurol. 65, 618 (2009).

64. Berardelli, A., Rothwell, J. C., Thompson, P. D. & Hallett, M. Pathophysiology of bradykinesia in Parkinson’s disease. Brain 124, 2131–2146 (2001).

65. Braak, H. et al. Staging of brain pathology related to sporadic Parkinson’s disease. Neurobiol Aging 24, 197–211 (2003).

66. Riemensperger, T. et al. A single dopamine pathway underlies progressive locomotor deficits in a Drosophila model of Parkinson disease. Cell Rep 5, 952–960 (2013).

67. Varga, S. J., Qi, C., Podolsky, E. & Lee, D. A new Drosophila model to study the interaction between genetic and environmental factors in Parkinson’s disease. Brain Res 1583, 277–286 (2014).

68. Gerber, B. & Stocker, R. F. The Drosophila larva as a model for studying chemosensation and chemosensory learning: a review. Chem Senses 32, 65–89 (2007).

69. Gerber, B., Biernacki, R. & Thum, J. Odor-taste learning assays in Drosophila larvae. Cold Spring Harb Protoc 2013, 212–223 (2013).

70. Inagaki, H. K. et al. Optogenetic control of Drosophila using a red-shifted channelrhodopsin reveals experience-dependent influences on courtship. Nat Methods 11, 325–32 (2014).

71. Boto, T., Stahl, A. & Tomchik, S. M. Cellular and circuit mechanisms of olfactory associative learning in Drosophila. https://doi.org/10.1080/01677063.2020.1715971 34, 36–46 (2020).

72. King, A. N. et al. A Peptidergic Circuit Links the Circadian Clock to Locomotor Activity. Current Biology 27, 1915–1927.e5 (2017).

73. Shafer, O. T. et al. Widespread receptivity to neuropeptide PDF throughout the neuronal circadian clock network of Drosophila revealed by real-time cyclic AMP imaging. Neuron 58, 223–37 (2008).

74. Fernandez-Chiappe, F. et al. Dopamine Signaling in Wake-Promoting Clock Neurons Is Not Required for the Normal Regulation of Sleep in Drosophila. The Journal of Neuroscience 40, 9617–9633 (2020).

75. Pfeiffenberger, C., Lear, B. C., Keegan, K. P. & Allada, R. Locomotor Activity Level Monitoring Using the Drosophila Activity Monitoring (DAM) System. Cold Spring Harb Protoc 2010, pdb.prot5518 (2010).

76. Aso, Y. et al. Mushroom body output neurons encode valence and guide memory- based action selection in Drosophila. Elife 3, e04580 (2014).

77. Moreira, J.-M. et al. optoPAD, a closed-loop optogenetics system to study the circuit basis of feeding behaviors. Elife 8, (2019).

78. Koyanagi, M. & Terakita, A. Diversity of animal opsin-based pigments and their optogenetic potential. Biochimica et Biophysica Acta (BBA) - Bioenergetics 1837, 710– 716 (2014).

79. Muntean, B. S. et al. Interrogating the Spatiotemporal Landscape of Neuromodulatory GPCR Signaling by Real-Time Imaging of cAMP in Intact Neurons and Circuits. Cell Rep 22, 255–268 (2018).

80. Lobingier, B. T. & von Zastrow, M. When trafficking and signaling mix: How subcellular location shapes G protein-coupled receptor activation of heterotrimeric G proteins. Traffic 20, 130–136 (2019).

81. Anton, S. E. et al. Receptor-associated independent cAMP nanodomains mediate spatiotemporal specificity of GPCR signaling. Cell 185, 1130–1142.e11 (2022).

82. Spoida, K. et al. Melanopsin Variants as Intrinsic Optogenetic on and off Switches for Transient versus Sustained Activation of G Protein Pathways. Current Biology 26, 1206–1212 (2016).

83. Koyanagi, M. et al. High-performance optical control of GPCR signaling by bistable animal opsins MosOpn3 and LamPP in a molecular property-dependent manner. Proc Natl Acad Sci U S A 119, e2204341119 (2022).

84. Edgar, R. C. MUSCLE: a multiple sequence alignment method with reduced time and space complexity. BMC Bioinformatics 2004 5:1 5, 1–19 (2004).

85. Han, C., Jan, L. Y. & Jan, Y.-N. Enhancer-driven membrane markers for analysis of nonautonomous mechanisms reveal neuron-glia interactions in Drosophila. Proc Natl Acad Sci U S A 108, 9673–8 (2011).

86. Tully, T., Preat, T., Boynton, S. C. & Del Vecchio, M. Genetic dissection of consolidated memory in Drosophila. Cell 79, 35–47 (1994).

87. Pfeiffer, B. D. et al. Refinement of tools for targeted gene expression in Drosophila. Genetics 186, 735–55 (2010).

88. Pielage, J., Bulat, V., Zuchero, J. B., Fetter, R. D. & Davis, G. W. Hts/Adducin controls synaptic elaboration and elimination. Neuron 69, 1114–31 (2011).

89. Nern, A., Pfeiffer, B. D. & Rubin, G. M. Optimized tools for multicolor stochastic labeling reveal diverse stereotyped cell arrangements in the fly visual system. Proceedings of the National Academy of Sciences 201506763 (2015) doi:10.1073/pnas.1506763112.

90. Schmid, B., Helfrich-Förster, C. & Yoshii, T. A new ImageJ plug-in ‘actogramJ’ for chronobiological analyses. J Biol Rhythms 26, 464–467 (2011).

91. Schlichting, M. & Helfrich-Förster, C. Photic Entrainment in Drosophila Assessed by Locomotor Activity Recordings. Methods Enzymol 552, 105–123 (2015).

92. Kobler, J. M., Rodriguez Jimenez, F. J., Petcu, I. & Grunwald Kadow, I. C. Immune Receptor Signaling and the Mushroom Body Mediate Post-ingestion Pathogen Avoidance. Curr Biol 30, 4693–4709.e3 (2020).

