## Supplemental figures and table for "Optimized design and *in vivo* application of optogenetically functionalized *Drosophila* dopamine receptors"

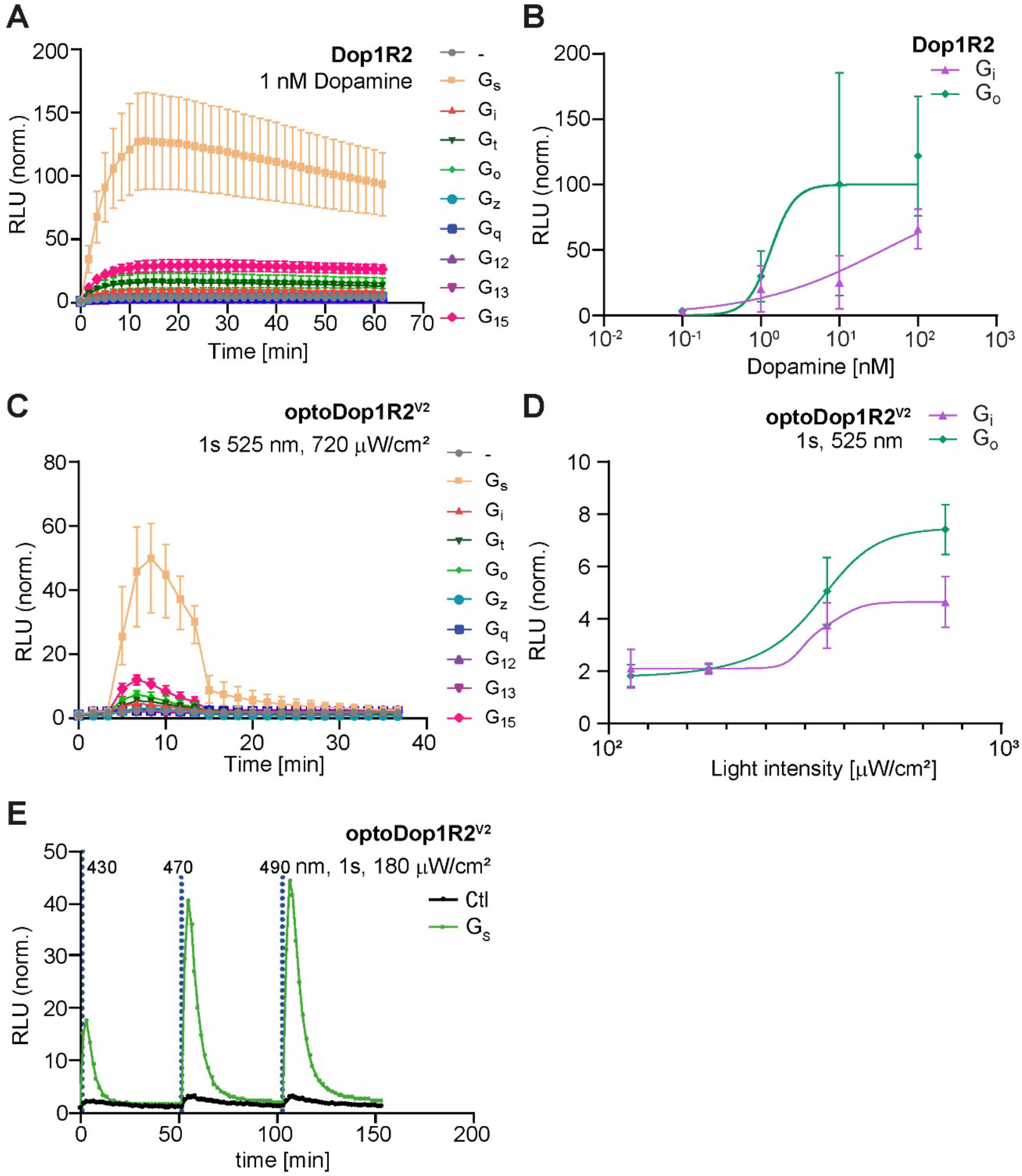


**Figure S1, related to Figure 1: Validation of optoDop1R1$^{V2}$**

(A) Kinetic G protein coupling properties of *Drosophila* Dop1R2 with 1nM DA (n=4). (B) DA concentration dependent maximum activation of $G_s$ and $G_{15}$ signaling of Dop1R1 (n=4). (C) G protein coupling properties of optoDop1R1$^{V1}$ after activation with light (1s 525 nm, 720 µW/cm$^2$). Normalized response kinetics are shown (n=7). (D) G protein coupling properties of improved optoDop1R1$^{V2}$ after activation with light (1s 525 nm, 720 µW/cm$^2$). Maximum normalized response kinetics are shown (n=7). (E) Wavelength-dependent induction of $G_s$-mediated cAMP production after optoDop1R1$^{V2}$ activation with light (1s 180 µW/cm$^2$, 430-490 nm, n=3).

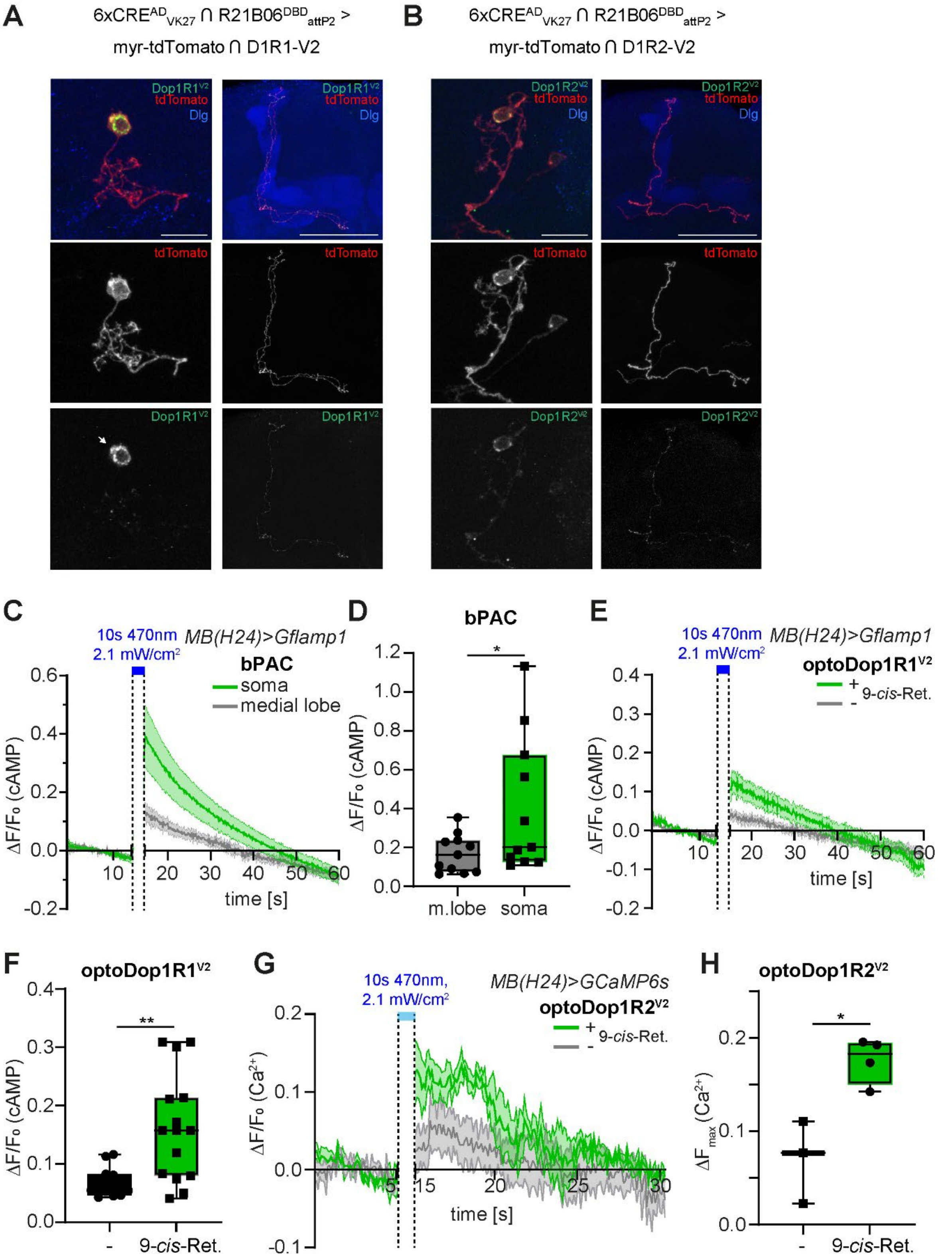


**Figure S2, related to Figure 2: Validation of optoDop1R2$^{V2}$**

(A) Kinetic G protein coupling properties of Drosophila Dop1R1 with 1nM DA (n=4). (B) DA concentration dependent maximum activation of $G_i$ and $G_o$ signaling of Dop1R2 (n=3-4). (C) G protein coupling properties of optoDop1R2$^{V2}$ after activation with light (1s 525 nm, 720 μW/cm$^2$). Normalized response kinetics are shown (n=4). (D) Light intensity-dependent maximum of $G_i$ and $G_o$ signaling induced by optoDop1R2$^{V2}$ (1s 525 nm, n=3-4). (E) Wavelength-dependent induction of $G_s$-mediated cAMP production after optoDop1R1$^{V2}$ activation with light (1s 180 μW/cm$^2$, 430-490 nm, n=3).

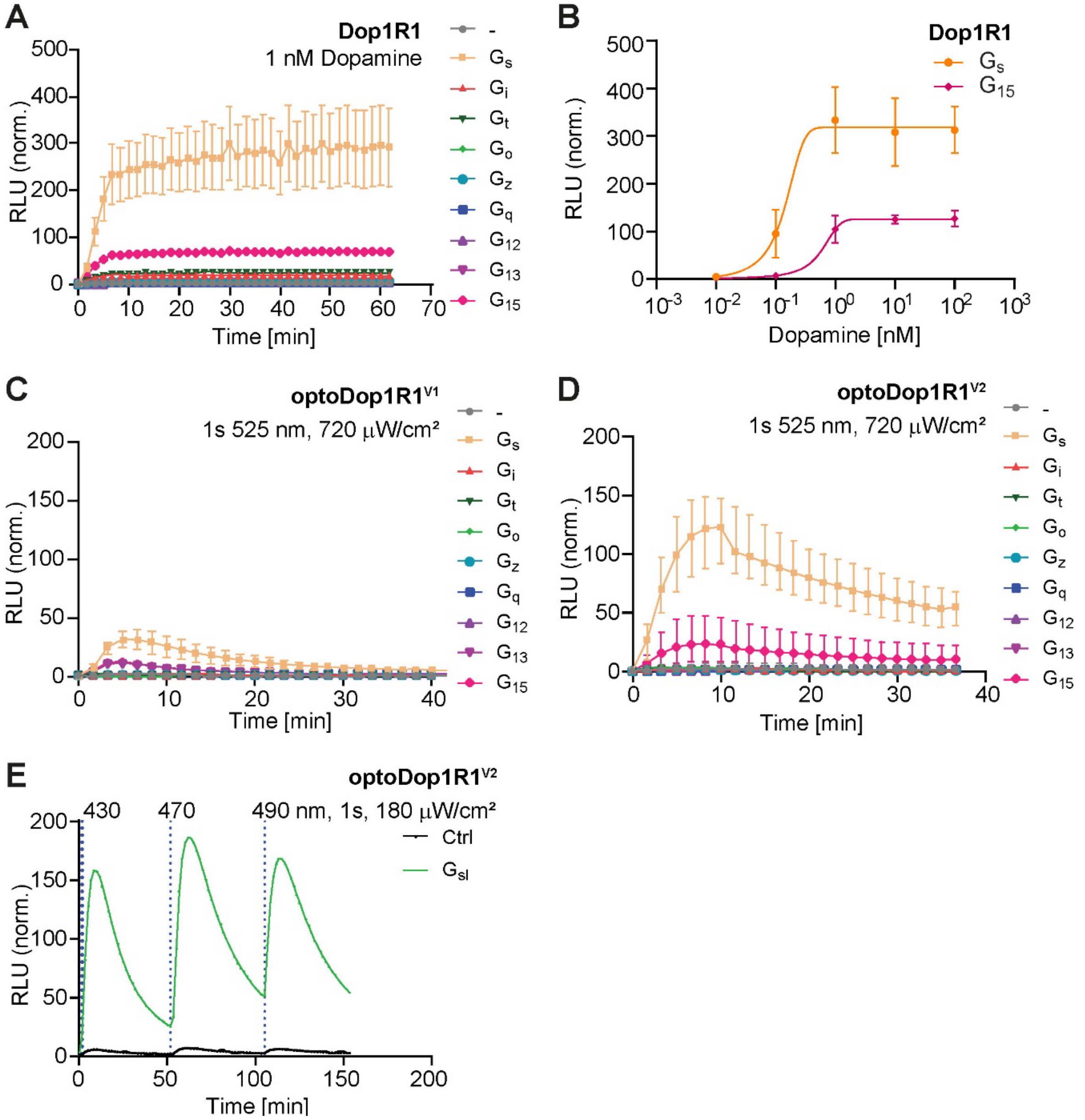


**Figure S3, related to Figure 3: *In vivo* localization and validation of optoDopR activity**
(A) Single cell expression of optoDop1R1$^{V2}$ in the adult mushroom body using activity dependent induction of Gal4 activity [52]. Examples of KCs labeled with tdTomato expressing optoDop1R1$^{V2}$ (scale bar 10 µm, 50 µm). Expression can be detected throughout the KC axon with punctate patterns. (B) Single cell expression of optoDop1R2$^{V2}$ in the adult mushroom body with examples of KCs labeled with tdTomato expressing optoDop1R2$^{V2}$ (scale bar 10 µm, 50 µm). Prominent axonal localization can be detected along the KC axons. (C) cAMP responses in the larval mushroom body monitored using Gflamp1 and induced by bPAC activation with blue light (*H24-Gal4>G-Flamp1, bPAC*, 10s 470 nm, n=11). Responses in the soma (green) and medial lobe (grey) are shown over time. (D) Maximum cAMP responses in the KC soma and MB medial lobe after light-induced activation of bPAC (10s 470 nm, n=11, 11). (E) cAMP imaging in the larval mushroom body using Gflamp1 and optoDop1R1$^{V2}$ expression (*H24-Gal4>G-Flamp1, optoDop1R1$^{V2}$*, 10s 470 nm, n=11, 15). Responses in the soma after 10s blue light illumination are shown over time. (F) Maximum cAMP responses in the KC somata after light-induced activation of optoDop1R1$^{V2}$ (10s 470 nm, n=11, 15). (G) *In vivo* calcium imaging of optoDop1R2$^{V2}$ expressed in the larval mushroom body using GCaMP6s. Neuronal calcium responses in KC somata in animals reared with or without 9-*cis*-retinal are shown over time (*H24-Gal4>GCaMP6s, optoDop1R2$^{V2}$*, 10s 470 nm, n=5, 5). (H) Maximum responses in the KC somata after light-induced activation of optoDop1R2$^{V2}$ (10s 470 nm, n=3, 4).

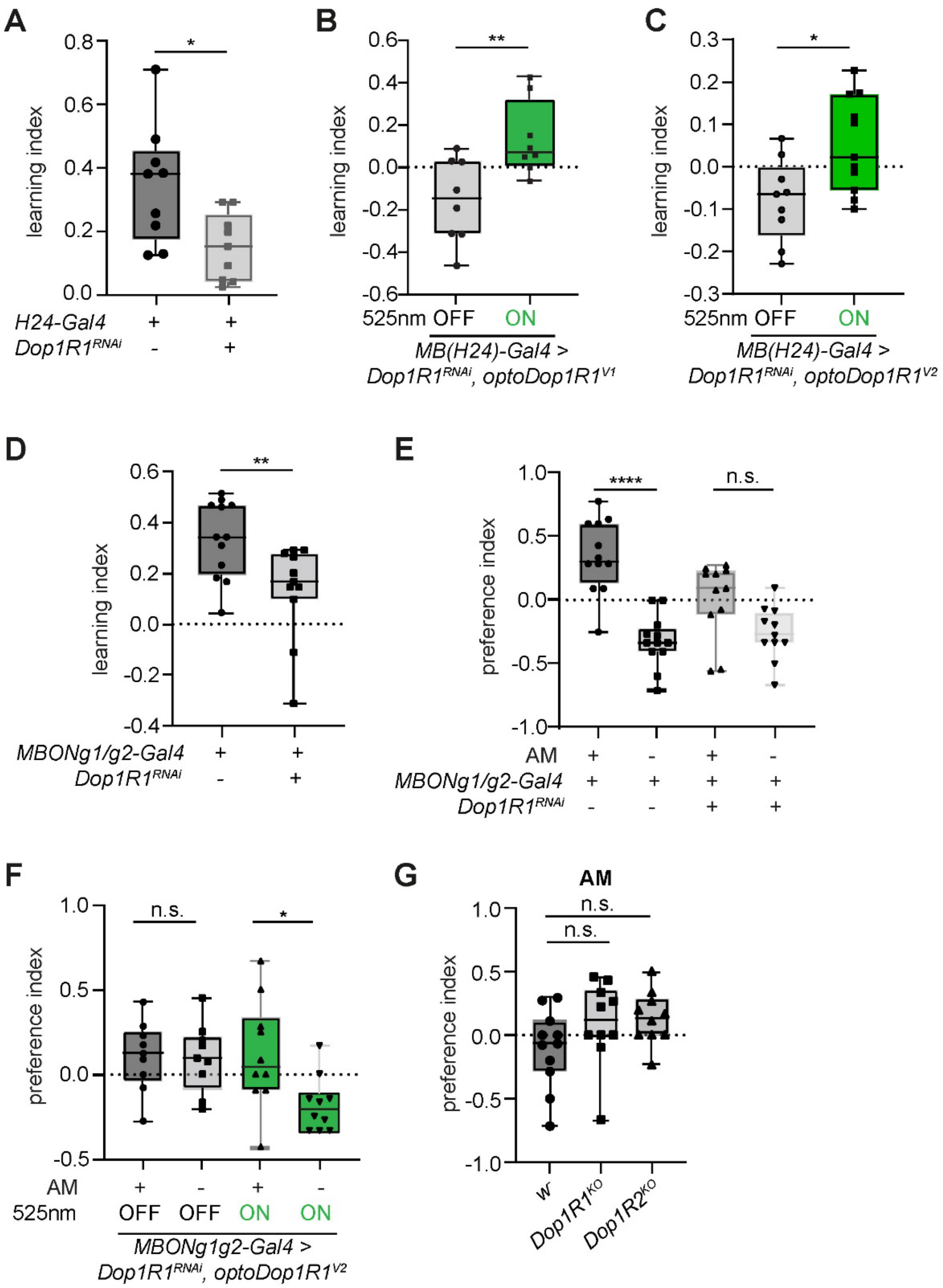


**Figure S4, related to Figure 4: Functional validation of optoXRs in Drosophila larvae *in vivo***
(A) Larval learning is impaired upon Dop1R1$^{RNAi}$ expression in the MB (*H24-Gal4> Dop1R1$^{RNAi}$*, n=9, 9). (B) Dop1R1-dependent single odor-fructose learning in larvae. Animals expressing optoDop1R1$^{V1}$ and Dop1R1$^{RNAi}$ in KCs were trained using fructose-odor learning (3x3min) with or without light activation during fructose exposure (*H24-Gal4>optoDop1R1$^{V1}$*, 3 min 525 nm, 720 µW/cm$^2$). Learning index of 9-*cis*-retinal fed animals with and without light activation during training are shown (n=8, 8, *p<0.05). (C) Single odor-fructose learning in larvae expressing optoDop1R1$^{V2}$ and Dop1R1$^{RNAi}$ in KCs. Fructose-odor learning (3x3min) with or without light activation during fructose exposure (*H24-Gal4>optoDop1R1$^{V2}$*, 3 min 525 nm, 720 µW/cm$^2$). Learning index of 9-*cis*-retinal fed animals with and without light activation during training are shown (n=9, 11, *p<0.05). (D) Larval learning is impaired upon Dop1R1$^{RNAi}$ expression in the MBON-g1/g2 (*MBON-g1/g2>Dop1R1$^{RNAi}$*, n=12, 11). (E) Fructose reward learning-dependent induction of odor preference (AM or blank) for Dop1R1-dependent data from (D). (F) Learning-dependent induction of odor preference (AM or blank) for optoDop1R1-

dependent rescue of Dop1R1$^{RNAi}$ in MBON-g1/g2 from Fig. 4E. (G) Innate preference index for AM in control (w$^{-}$), *Dop1R1$^{KO}$* and *Dop1R2$^{KO}$* 3$^{rd}$ instar larvae (n=11, 10, 9, *p<0.05).

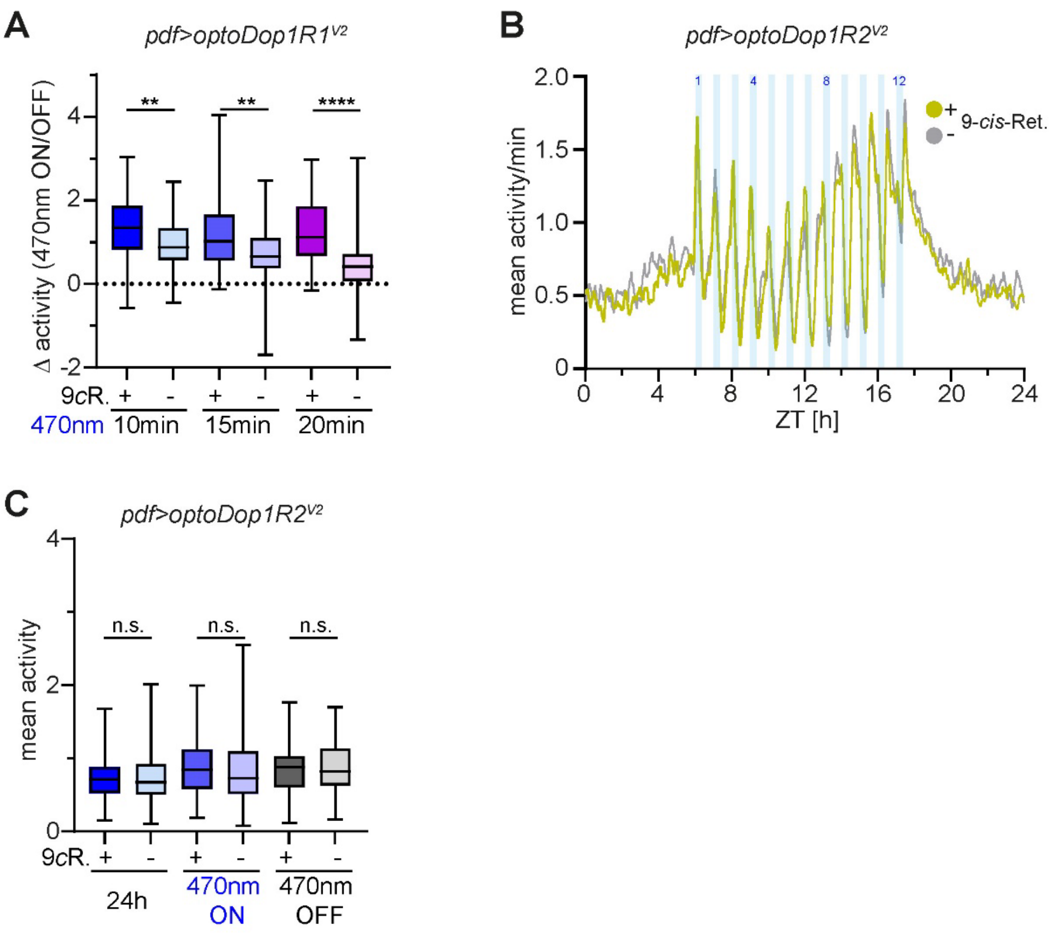


**Figure S5, related to Figure 5: Cell type-specific function of Dop1R1 activity in blue light induced arousal**

(A) Activity difference of flies expressing optoDop1R1$^{V2}$ in pdf neurons (with and without 9cR feeding) during light on times using different duration of blue light pulse exposure (1/h, 10, 15 or 20min, n=83/77, p<0.0001). (B) Mean activity during 24h monitoring in flies expressing optoDop1R2$^{V2}$ in pdf neurons with and without 9cR feeding (n=90). Blue light pulses (12x 20min, 1/h) during daytime increase fly activity independently of optoDop1R1$^{V2}$ activation. (C) Mean activity of pdf>optoDop1R2$^{V2}$ -expressing flies during the entire 24h, all light on and light off phases (n=90, p>0.05).

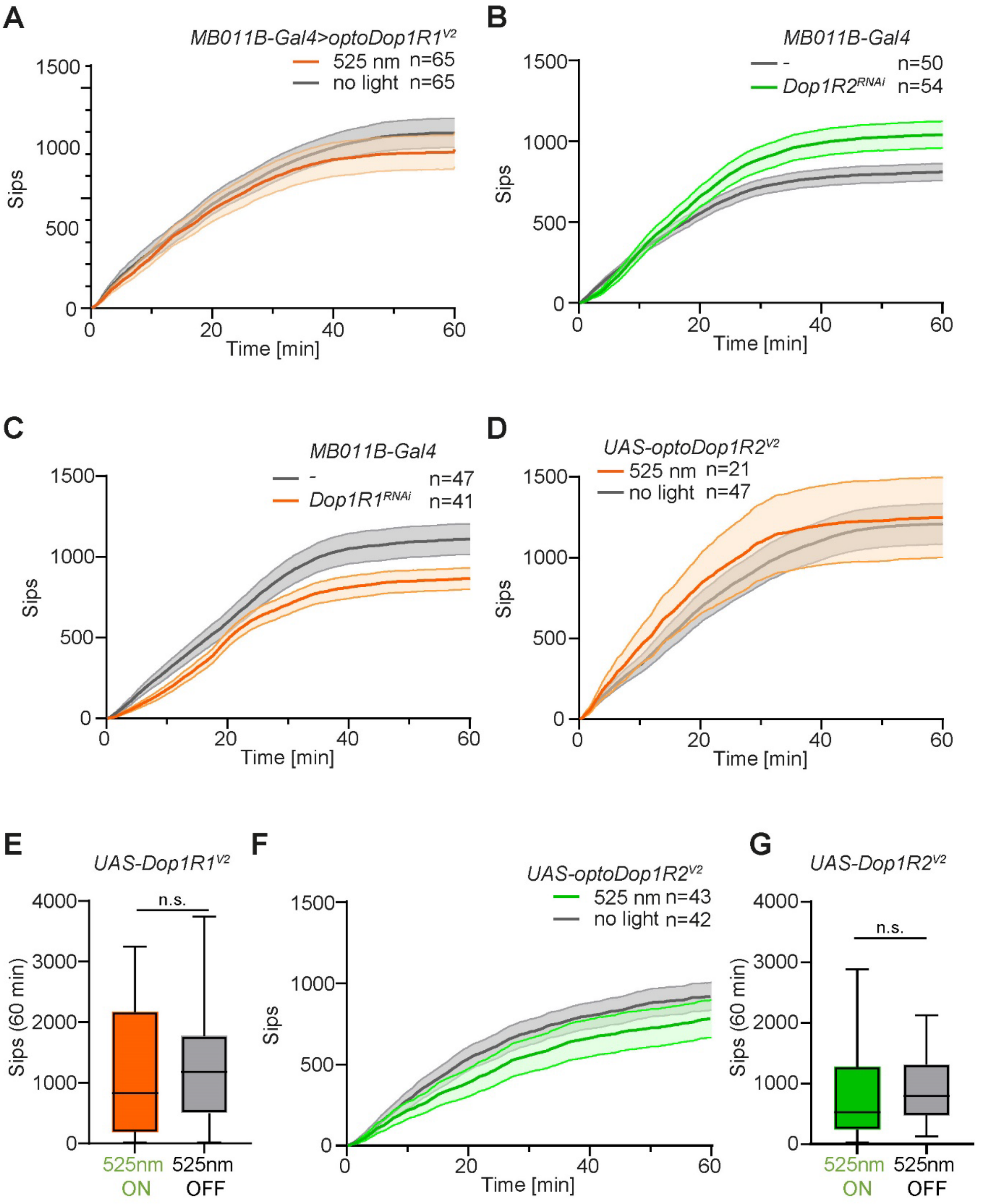


**Figure S6, related to Fig. 6: Cell type-specific function of Dop1R2 activity in satiety**
(A) Cumulative sips over time in flies expressing optoDop1R1$^{V2}$ with *MB011B-Gal4* with or without light stimulation (n=65,65). (B) Cumulative sips over time in flies expressing Dop1R2$^{RNAi}$ with *MB011B-Gal4* compared to control (n=50,54). (C) Cumulative sips over time in flies expressing Dop1R1$^{RNAi}$ with *MB011B-Gal4* (n=47,41). (D) Cumulative sips over time in optoDop1R1$^{V2}$ transgenes without Gal4 expression and with or without light stimulation (n=21,48). (E) Average sips at 60min for optoDop1R1$^{V2}$ transgenes without Gal4 expression and with or without light stimulation (n=21,48). (F) Cumulative sips over time in optoDop1R2$^{V2}$ transgenes without Gal4 expression and with or without light stimulation (n=43,42). (G) Average sips at 60min for optoDop1R2$^{V2}$ transgenes without Gal4 expression and with or without light stimulation (n=43,42).

**Table S1. Previous optoXRs and their in vivo applications.** Only chimeric optoXRs aiming to mimic target receptor signaling and function are shown. Abbreviations: Rho: bovine Rhodopsin, OPN4: melanopsin

| **Chimeric receptor** | **Original reference** | ***In vivo* applications** | **Cell type-specificity/rescue of endogenous receptor function** |
|---|---|---|---|
| Rho:$\beta_2$AR | (Kim et al. 2005) | - virus-mediated overexpression in mouse N. accumbens neurons (Airan et al. 2009)<br>- virus-mediated overexpression in mouse basolateral amygdala, promoting anxiety-like behavior (Siuda et al. 2015b, 2016) | partial/no<br>partial/no |
| Rho:$\alpha_1$AR | (Airan et al. 2009) | - virus-mediated overexpression in mouse N. accumbens neurons, reward-related preference behavior<br>- virus-mediated overexpression in mouse CA1 astrocytes, memory acquisition(Adamsky et al. 2018)<br>- transgenic overexpression in mouse cortical astrocytes, remote memory acquisition (Iwai et al. 2021)<br>- virus-mediated overexpression in mouse astrocytes in slices, electrophysiology (Gerasimov et al. 2021) | partial/no<br>partial/no<br>partial/no<br>partial/no |
| Rho:µOR | (Barish et al. 2013) | - virus-mediated overexpression in mouse dorsal root ganglion neurons, preference/aversion behavior (Siuda et al. 2015a)<br>- Penk-Cre dependent virus-mediated overexpression in dorsal raphe nucleus subset neurons, restoration of consumption behavior (Castro et al. 2021) | partial/no<br>yes/yes |
| Rho:DRD1 | (Gunaydin et al. 2014) | - DRD1-Cre dependent virus-mediated overexpression in mouse N. accumbens; activation of medium spiny neurons to increase social interaction | yes/no |
| Rho:CXCR4 | (Xu et al. 2014) | - virus-mediated overexpression in mouse, T-cell recruitment | yes/no |
| Rho:$A_{2A}$R | (Li et al. 2015) | - virus-mediated overexpression in mouse hippocampus and N. accumbens, spatial memory performance and locomotor activity<br>- adora2a-cre dependent virus-mediated overexpression in mouse striatopallidal neurons, goal-directed behavior (Li et al. 2016) | partial/no<br>yes/no |
| OPN4:mGluR$_6$ | (van Wyk et al. 2015) | - virus-mediated overexpression in retinal ganglion cells, restoration of visually guided behavior<br>- virus-mediated overexpression in bipolar cells, restoration of visually guided behavior (Kralik et al. 2022) | yes/partial (degeneration model)<br>yes/yes (degeneration model) |
| Rho:Fz7 | (Čapek et al. 2019) | - Zebrafish mRNA injection and overexpression, mesoderm cell migration | no/yes |

### Supplemantary References

Adamsky A, Kol A, Kreisel T, Doron A, Ozeri-Engelhard N, et al. 2018. Astrocytic Activation Generates De Novo Neuronal Potentiation and Memory Enhancement. *Cell*. 174(1):59-71.e14

Airan RD, Thompson KR, Fenno LE, Bernstein H, Deisseroth K. 2009. Temporally precise in vivo control of intracellular signalling. *Nature*. 458(7241):1025–29

Barish PA, Xu Y, Li J, Sun J, Jarajapu YPR, Ogle WO. 2013. Design and functional evaluation of an optically active μ-opioid receptor. *Eur J Pharmacol*. 705(1–3):42–48

Čapek D, Smutny M, Tichy AM, Morri M, Janovjak H, Heisenberg CP. 2019. Light-activated Frizzled7 reveals a permissive role of non-canonical wnt signaling in mesendoderm cell migration. *Elife*. 8:

Castro DC, Oswell CS, Zhang ET, Pedersen CE, Piantadosi SC, et al. 2021. An endogenous opioid circuit determines state-dependent reward consumption. *Nature 2021 598:7882*. 598(7882):646–51

Gerasimov E, Erofeev A, Borodinova A, Bolshakova A, Balaban P, et al. 2021. Optogenetic Activation of Astrocytes—Effects on Neuronal Network Function. *International Journal of Molecular Sciences 2021, Vol. 22, Page 9613*. 22(17):9613

Gunaydin LA, Grosenick L, Finkelstein JC, Kauvar I V., Fenno LE, et al. 2014. Natural Neural Projection Dynamics Underlying Social Behavior. *Cell*. 157(7):1535–51

Iwai Y, Ozawa K, Yahagi K, Mishima T, Akther S, et al. 2021. Transient Astrocytic Gq Signaling Underlies Remote Memory Enhancement. *Front Neural Circuits*. 15:

Kim J-M, Hwa J, Garriga P, Reeves PJ, RajBhandary UL, Khorana HG. 2005. Light-Driven Activation of β 2 -Adrenergic Receptor Signaling by a Chimeric Rhodopsin Containing the β 2 -Adrenergic Receptor Cytoplasmic Loops. *Biochemistry*. 44(7):2284–92

Kralik J, van Wyk M, Stocker N, Kleinlogel S. 2022. Bipolar cell targeted optogenetic gene therapy restores parallel retinal signaling and high-level vision in the degenerated retina. *Communications Biology 2022 5:1*. 5(1):1–15

Li P, Rial D, Canas PM, Yoo JH, Li W, et al. 2015. Optogenetic activation of intracellular adenosine A2A receptor signaling in the hippocampus is sufficient to trigger CREB phosphorylation and impair memory. *Mol Psychiatry*. 20(11):1481

Li Y, He Y, Chen M, Pu Z, Chen L, et al. 2016. Optogenetic Activation of Adenosine A2A Receptor Signaling in the Dorsomedial Striatopallidal Neurons Suppresses Goal-Directed Behavior. *Neuropsychopharmacology*. 41(4):1003–13

Oh E, Maejima T, Liu C, Deneris E, Herlitze S. 2010. Substitution of 5-HT1A Receptor Signaling by a Light-activated G Protein-coupled Receptor. *Journal of Biological Chemistry*. 285(40):30825–36

Siuda ER, Al-Hasani R, McCall JG, Bhatti DL, Bruchas MR. 2016. Chemogenetic and Optogenetic Activation of Gαs Signaling in the Basolateral Amygdala Induces Acute and Social Anxiety-Like States. *Neuropsychopharmacology*. 41(8):2011–23

Siuda ER, Copits BA, Schmidt MJ, Baird MA, Al-Hasani R, et al. 2015a. Spatiotemporal Control of Opioid Signaling and Behavior. *Neuron*. 86(4):923–35

Siuda ER, McCall JG, Al-Hasani R, Shin G, Il Park S, et al. 2015b. Optodynamic simulation of β-adrenergic receptor signalling. *Nat Commun*. 6(1):8480

van Wyk M, Pielecka-Fortuna J, Löwel S, Kleinlogel S. 2015. Restoring the ON Switch in Blind Retinas: Opto-mGluR6, a Next-Generation, Cell-Tailored Optogenetic Tool. *PLoS Biol*. 13(5):1–30

Xu Y, Hyun YM, Lim K, Lee H, Cummings RJ, et al. 2014. Optogenetic control of chemokine receptor signal and T-cell migration. *Proc Natl Acad Sci U S A*. 111(17):6371–76

**Supplementary Movies**

**Movie S1**

Blue light induced rolling of larvae expressing bPAC in nociceptors (*ppk-Gal4>UAS-bPAC*)

**Movie S2**

Blue light induced rolling of larvae expressing optoDop1R1$^{V1}$ in nociceptors (*ppk-Gal4>UAS-optoDop1R1*$^{V1}$)

**Movie S3**

Blue light induced rolling of larvae expressing optoDop1R1$^{V2}$ in nociceptors (*ppk-Gal4>UAS-optoDop1R1*$^{V2}$)

**Movie S4**

Blue light induced rolling of larvae expressing optoDop1R2$^{V2}$ in nociceptors (*ppk-Gal4>UAS-optoDop1R2*$^{V2}$)

**Movie S5**

Blue light induced rolling of larvae expressing CsChrimson in nociceptors (*ppk-Gal4>UAS-CsChrimson*)

**Movie S6**

cAMP response of blue light induced bPAC activation in the mushroom body medial lobe (*H24-Gal4>UAS-bPAC,UAS-Gflamp1*)

**Movie S7**

cAMP response of blue light induced optoDop1R1$^{V2}$ activation in the mushroom body medial lobe (*H24-Gal4>UAS-optoDop1R1*$^{V2}$*,UAS-Gflamp1*)

**Movie S8**

Locomotion of rotenone-treated larvae expressing optoDop1R1$^{V2}$ in the endogenous pattern of Dop1R1 using a knock-in Gal4 line (*Dop1R1*$^{KO\text{-}Gal4}$*>UAS-optoDop1R1*$^{V2}$). Larvae were tracked in the dark (magenta tracks) and subsequently upon green light illumination (green tracks).
